# A stop-gained mutation in a Peroxiredoxin gene may underlie the low-chill phenotype of the apricot spontaneous mutant ‘Búlida Precoz’

**DOI:** 10.1101/2025.07.30.667652

**Authors:** Daniel González-Palazón, David Ruiz, José A. Campoy Corbalán

## Abstract

Somatic mutations in clonally propagated fruit trees, known as “bud sports,” are a primary source of new and agronomically valuable traits. This study addresses the study of ‘Búlida Precoz’, a spontaneous somatic variant of the ‘Búlida’ apricot characterized by an early flowering (low-chill) phenotype. The main objective of this work was to design and implement a comprehensive bioinformatic workflow to identify the specific somatic mutation(s) responsible for this significant developmental change, using whole-genome sequencing (WGS) data.

A comparative genomics approach was employed, analyzing the genomes of four biological replicates from the ‘Búlida Precoz’ mutant and four from the original ‘Búlida’ wild-type. The analysis was centered on a specialized somatic variant-calling workflow using `GATK Mutect2`, chosen for its high sensitivity in detecting low-frequency mutations. The results were cross-validated against an independent analysis using the germline caller `GATK HaplotypeCaller.`

The final candidates were functionally annotated to predict their biological impact on gene function. This work establishes a solid foundation for future experimental validation and provides valuable molecular targets that could be used in marker-assisted breeding programs aimed at developing new apricot cultivars better adapted to climates with mild winters.

## 2. INTRODUCTION

Climate change is exerting considerable pressure on global agricultural systems, with temperate fruit production being particularly vulnerable to the impacts of rising temperatures. A critical aspect of this vulnerability lies in the disruption of dormancy cycles in temperate fruit trees like apricot (*Prunus armeniaca* L.). These species have evolved a chilling requirement, a specific period of exposure to low temperatures, to ensure bud break and flowering are synchronized with favorable spring conditions, thereby mitigating the risk of frost damage to reproductive structures and securing fruit set. However, the progressive increase in global mean temperatures is leading to milder winters, frequently resulting in insufficient chill accumulation. This directly compromises flowering synchrony, fruit development, and overall orchard productivity for many established cultivars (Ruiz et al., 2019). Addressing the challenge of adapting fruit crops to warming climates through traditional breeding for low-chill requirement cultivars is typically a challenging and lengthy process, often constrained by the limited availability of low-chill requirement trait donors. In this scenario, spontaneous somatic mutations that manifest as “bud sports” represent a valuable resource for achieving rapid crop adaptation. These naturally, yet improbable, occurring genetic changes within an individual branch meristematic layers can lead to the emergence of novel and agronomically desirable phenotypes, such as modified chilling requirements or altered flowering times, directly within an established and commercially successful cultivar. The clonal propagation of such bud sports allows for the direct fixation and utilization of these traits, bypassing generations, and decades, of conventional breeding (Goel et al., 2024; Ruiz et al., 2019; Sun et al., 2024). A notable example is the spontaneous mutant “Búlida Precoz,” discovered in southern Spain and derived from the commercial apricot cultivar “Búlida.” This mutant displays significantly earlier flowering times compared to its wild-type counterpart, indicating a reduced chilling requirement. Phenotypic analyses have shown that “Búlida Precoz” accumulates significantly fewer chill portions (CP) 33.7 compared to the “Búlida” wild type (47.5 CP), resulting in flowering approximately 17 days earlier under the same environmental conditions (Ruiz et al., 2019). Similarly, natural mutants like “Rojo Pasión Precoz” and “Santa Rosa Precoz” have also demonstrated substantial reductions in chilling requirements and advances in flowering time, confirming the stability and agronomic value of such mutations across multiple growing seasons. Similar findings for other natural mutants like ‘Rojo Pasión Precoz’ (apricot) and ‘Santa Rosa Precoz’ (Japanese plum) have also demonstrated substantial reductions in chilling requirements (8.0 and 13.1 Chill Portions, respectively) and advancements in flowering time, further underscoring the potential of such spontaneous sports for horticultural improvement (Ruiz et al., 2019). Understanding the genetic basis underlying these spontaneous mutations can significantly advance breeding programs aimed at adapting fruit trees to warmer climates. Previous studies have revealed that the majority of somatic mutations in plants occur in layer-specific patterns within the meristematic regions (L1, L2, L3), demonstrated in the ‘Rojo Pasión’ apricot cultivar (derived from ‘Orange Red’ and ‘Currot’) that over 90% of somatic mutations are confined to individual meristematic layers. This is of great importance for somatic mutation identification: departing form the assumption of an equally distributed number of cells per layer (although we know that L1 is usually less represented), one heterozygous mutation would be present in just 1/6^th^ of the reads (1/ 3 layers ^*^ 1/ 2 genome copies in diploid = 1/6).

This theoretical model, where a heterozygous mutation in a single meristematic layer is present in approximately 1/6 of the total alleles (assuming even distribution of cells among the three layers), would result in an expected Variant Allele Frequency (VAF) of around 16.7% (**Figure 1**). The detection of such a low-frequency signal presents a challenge, requiring high sequencing depth to be confidently distinguished from background sequencing errors.

**Figure 1.**
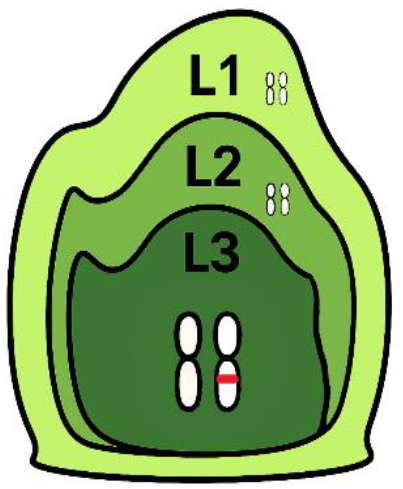
Theoretical Model of a Layer-Specific Somatic Mutation. The diagram illustrates a heterozygous mutation (in red) occurring in one of the three diploid meristematic layers (L3). This shows how, assuming even distribution of cells among the three layers, the mutation would be present in only 1/6 of the total alleles within a bulk tissue sample, leading to a low expected allele frequency (ca. 1/6th). Chromosomes in L3 are bigger for illustration purposes.

It was also revealed that the L1 (epidermis precursor) showed a higher mutation load compared to L2 (mesophyll and germline precursor) and confirmed that these mutations are propagated clonally through axillary meristems, following the tree’s branching architecture. The generation of a haplotype-resolved genome assembly for ‘Rojo Pasión’, achieved using PacBio HiFi sequencing and a gamete-binning approach, has provided a critical resource for genomic research and breeding in apricot (Campoy et al., 2020). Furthermore, single-cell RNA sequencing analyses have validated that these layer-specific mutations are transcribed exclusively within cells originating from the respective layers, thus establishing a direct link between the genetic alteration and its potential functional impact at the cellular level (Goel et al., 2024).

Complementary research on other fruit tree sports, such as the ‘RubyMac’ apple, has further illuminated the landscape of somatic genetic changes. A recent work focused on the ‘RubyMac’ sport, which exhibits altered fruit coloration, identified not only de novo point mutations (Gain-of-Heterozygosity, or GOH events) but also a significant number of somatic gene conversions leading to Loss-of-Heterozygosity (LOH) that distinguished the mutant branches from the wild-type portions of the tree.

A key finding of that study was that a notable fraction of these somatic variants, including a specific set of *de novo* mutations identified within individual haplotypes (haplotigs), exhibited allele frequencies consistent with a layer-specific origin, providing strong evidence for the prevalence of such mosaicism in bud sports. These studies also highlighted distinct mutational spectra; for instance, de novo mutations were often enriched for GC to AT transitions, whereas gene conversions showed a bias towards AT to GC transitions, suggesting a potential genomic mechanism that might counteract de novo mutational biases and influence GC content (Sun et al., 2024).

This study directly builds upon this background, primarily aiming to identify candidate causal somatic mutations for the early flowering phenotype observed in the ‘Búlida Precoz’ apricot mutant through a comparative analysis of its whole-genome resequencing data against that of the ‘Búlida’ wild-type. The methodology will involve a comprehensive bioinformatic analysis of Illumina paired-end sequencing data derived from several branches of both the mutant and wild-type trees. It is anticipated that the findings will enhance the understanding of flowering time regulation in apricot and potentially identify molecular markers beneficial for breeding programs focused on climate change adaptation.

## 3. MATERIAL AND METHODS

This section outlines the bioinformatic procedures employed for processing and analyzing the whole-genome resequencing data from ‘Búlida’ wild-type and ‘Búlida Precoz’ mutant apricot samples. The overall workflow is illustrated in **Figure 2**. Each step of the analysis was automated through a series of command-line scripts, which are fully documented and publicly available for reproducibility in the project’s GitHub repository.

**Figure 2.**
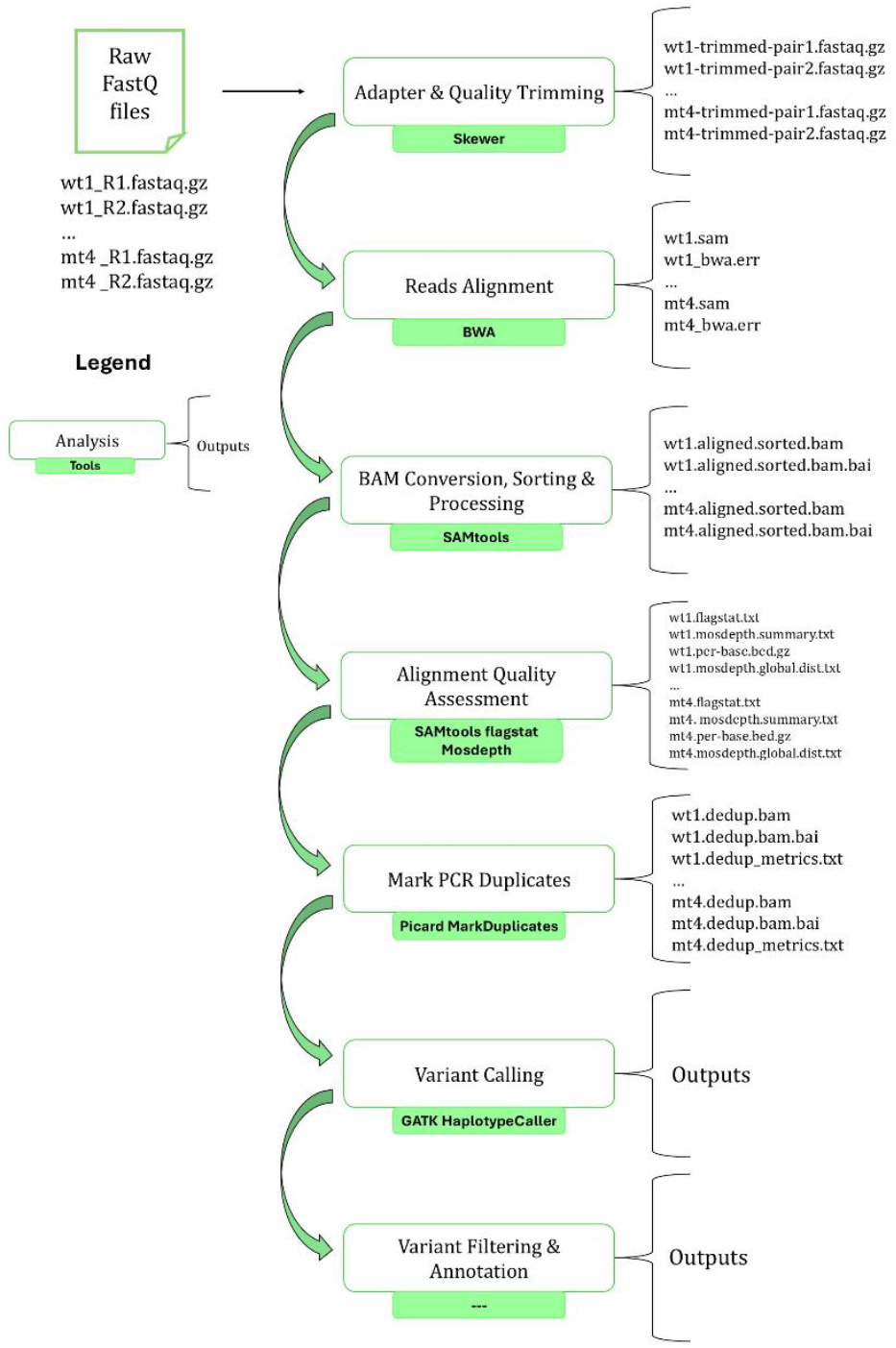
Overview of the bioinformatic pipeline for processing Illumina whole-genome resequencing data and identifying somatic variants. The pipeline starts with raw FASTQ files from ‘Búlida’ wild-type (wt1-wt4) and ‘Búlida Precoz’ mutant (mt1-mt4) samples. Key stages include: (1) Initial Quality Control using FastQC; (2) Adapter and Quality Trimming using Skewer; (3) Reads Alignment to the reference genome using BWA-MEM; (4) BAM Conversion, Sorting & Processing using SAMtools (including name sort, fixmate, coordinate sort, and indexing); (5) Alignment Quality Assessment using SAMtools flagstat and Mosdepth; (6) Marking of PCR Duplicates using Picard MarkDuplicates; and planned subsequent steps of (7) Variant Calling with GATK HaplotypeCaller and GATK Mutect2 (8) Variant Filtering & Annotation.

### 3.1. Plant Material and Sequencing Data

The plant material used in this study comprises the apricot (*Prunus armeniaca* L.) cultivar ‘Búlida’, serving as the wild-type (wt) control, and its spontaneous early-flowering mutant, ‘Búlida Precoz’ (mt). Both the wild-type and mutant genotypes were grafted onto commercial clonal rootstocks and cultivated under homogeneous orchard management conditions at the Experimental field of CEBAS-CSIC in Murcia, Spain, to minimize environmental variables. Only adult trees were used for analyses to avoid juvenility effects on flowering time and dormancy release.

For genomic analysis, DNA was extracted from leaves collected from four distinct branches of both the ‘Búlida’ wild-type and the ‘Búlida Precoz’ mutant trees to provide four biological replicates per genotype **(Figure 3)**. These samples represent bulk leaf tissue, encompassing all three meristematic cell layers (L1, L2, and L3). The samples were collected into Falcon tubes and directly snap-frozen in liquid nitrogen.

**Figure 3.**
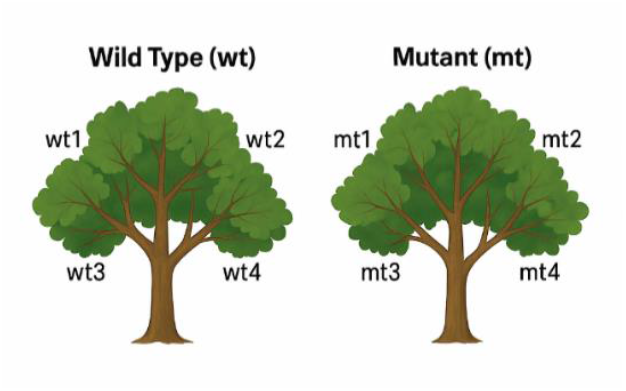
Schematic layout of sampling strategy. Fictitious depiction of a wild-type (wt) apricot tree (left) and a mutant (mt) apricot tree (right), each with four distinct branches selected for genomic analysis. Branches on the wild-type tree are labeled wt1, wt2, wt3, and wt4. Branches on the mutant tree are labeled mt1, mt2, mt3, and mt4.

The DNA was extracted with Machery Nagel Kit NucleoSpin PlantII. Elution with 40 μl PE elution buffer. After the first elution, it was eluted again with the eluate. For the quality of the DNA, the OD260/280 and OD260/280 were measured with NanoDrop. For the exact concentration, the DNA was measured with Qubit. The extracted DNA samples were sent to Max Planck-Genome Center in Cologne (MPGC, wt/mut1-2) or Novogene (wt/mut 3-4) for library preparation and sequencing using Illumina paired-end sequencing platforms.

The raw sequencing data for these samples were obtained from two distinct sequencing sources:

1. **Genome Centre in Cologne (**MPGC): Data from this source provided two biological replicates from ‘Búlida’ wild-type branches (**wt1, wt2**) and two from ‘Búlida Precoz’ mutant branches (**mt1, mt2**).
2. **Novogene:** Data from this project provided two additional biological replicates from ‘Búlida’ wild-type branches (**wt3, wt4**) and two from ‘Búlida Precoz’ mutant branches (**mt3, mt4**).

### 3.2. Raw Data and Sample Identity Quality Control

Prior to any data processing and analysis, two critical quality control (QC) procedures were performed. The first was to assess the quality of the raw sequencing reads, and the second was to verify the genetic identity and integrity of all eight experimental samples. These steps are essential to ensure the reliability of the entire downstream workflow.

Initial quality reports for all raw FASTQ files were generated using the **FastQC** tool (*Babraham Bioinformatics FastQC A Quality Control tool for High Throughput Sequence Data*, s. f.). This analysis provided a comprehensive overview of key metrics for each sample’s sequencing data, including per-base quality scores, GC content, sequence duplication levels, and potential adapter contamination. The resulting HTML reports were reviewed to confirm that the raw data met the necessary quality standards for further processing.

Subsequently, a genetic-level identity check was performed to confirm the relationship between all eight samples and to rule out potential sample swaps or duplications. Using `bcftools isec` (Danecek et al., 2021), every possible pair of samples was systematically compared based on their shared and unique variants from a high-quality, filtered VCF file. The purpose of this analysis was to validate that the biological replicates within each group (mutant and wild-type) were more genetically similar to each other than to samples from the opposing group, but different enough to discard, for example, possible sampling swap or resequencing of the same DNA, thus confirming the integrity of the experimental design.

### 3.3. Raw Read Pre-processing (Adapter and Quality Trimming)

Adapter sequences and low-quality bases were removed from the raw paired-end reads using `Skewer (v0.2.2)` (Jiang et al., 2014). The tool was run in paired-end mode (`-m pe`), trimming bases from the 3’ end to meet a minimum quality threshold of 20 (-q 20). Following trimming, reads shorter than 75 bp were discarded (`-l 75`). The resulting cleaned paired-end FASTQ files for each sample were used for all subsequent alignment steps.

### 3.4. Read Alignment to Reference Genome

Following pre-processing, the cleaned, paired-end reads for each of the eight samples (wt1-wt4 and mt1-mt4) were aligned to the reference genome of ‘Búlida’.

#### 3.4.1. Reference Genome Preparation of ‘Búlida’ cultivar

The primary reference genome used for this study was an *ad hoc* ‘Búlida’ reference-guided assembly (developed by José A. Campoy at Schneeberger’s lab at MPIPZ, unpublished results), designated `BUL_cur_guided.v1.0.fasta`. This assembly was previously generated using PacBio and Illumina sequencing data from the ‘Búlida’ cultivar, with the ‘Rojo Pasión’ genome (Campoy et al., 2020; Goel et al., 2024). Prior to alignment, the reference genome was indexed using bwa index for compatibility with the BWA aligner (Li & Durbin, 2009). Additionally, a FASTA index file (.fai) was created using `samtools faidx` (Danecek et al., 2021) to allow random access to reference sequences, a requirement for downstream tools like `Picard` and `GATK`.

#### 3.4.2. Alignment Procedure

The trimmed paired-end FASTQ files for each of the eight samples were aligned to the indexed `BUL_cur_guided.v1.0.fasta` reference genome using the **BWA-MEM** algorithm (v0.7.17) (Li & Durbin, 2009), which is optimized for aligning longer Illumina reads. Key parameters were set to ensure compatibility with downstream processing tools.

- **Marking Secondary Alignments (-M):** This option was used to mark shorter, split-read alignment hits as secondary. This is crucial for compatibility with downstream tools, particularly `Picard MarkDuplicates` (*Picard Tools - By Broad Institute*, s. f.) and GATK (*Somatic Calling Is NOT Simply a Difference between Two Callsets*, 2024) variant callers.
- **Read Group Information (-R):** This information, specifying the sample name (SM), library (LB), and platform (PL), is essential for GATK tools to properly track data from different samples and libraries. The output from `BWA-MEM` for each sample was generated in SAM (Sequence Alignment Map) format and subsequently processed as described in the following section.

#### 3.4.3. Post-Alignment Data Processing

Following alignment with BWA-MEM, the resulting SAM files for each sample were processed into coordinate-sorted, indexed BAM (Binary Alignment Map) files using `SAMtools (v1.6)` (Danecek et al., 2021). This procedure first involved converting the text-based SAM files to the compressed binary BAM format (`samtools view`). The BAM files were then sorted by read name (`samtools sort -n`), a prerequisite for using `samtools fixmate` to ensure mate-pair information was consistent and complete. Following the mate-pair correction, the files were sorted again, this time by genomic coordinates. Finally, the coordinate-sorted BAM files were indexed using `samtools index` to create a .bai file, allowing for rapid random access to alignment data in specific genomic regions by downstream tools.

### 3.5. Alignment Quality Assessment

Following the alignment and BAM post-processing steps, a quality assessment was performed on the final coordinate-sorted and indexed BAM files for each of the eight samples. This assessment focused on evaluating key alignment metrics, such as the percentage of mapped and properly paired reads, and the mean depth and breadth of genome coverage, to ensure the data was suitable for downstream variant calling.

#### 3.5.1. Alignment Statistics with samtools flagstat

To obtain a quantitative summary of the alignment outcomes, `samtools flagstat` (Danecek et al., 2021) was utilized. This command computes various metrics based on the SAM flags of the aligned reads, including the total number of reads, the number and percentage of mapped and properly paired reads, the number of singleton reads, and reads whose mates mapped to different chromosomes.

#### 3.5.2. Coverage Analysis with mosdepth

The depth and breadth of sequencing coverage for each aligned BAM file were assessed using `mosdepth (v0.3.3)` (Pedersen & Quinlan, 2018). The analysis utilized the `BUL_cur_guided.v1.0.fasta` reference genome `(-f option`) to calculate per-base depth, mean coverage for the genome and each contig, and the fraction of the genome covered at various depth thresholds.

### 3.6. Duplicate Marking

PCR and optical duplicates in the coordinate-sorted BAM files were identified and flagged using `Picard MarkDuplicates (v3.4.0)` (*Picard Tools - By Broad Institute*, s. f.). Duplicates were flagged (`SAM flag 0×400`) rather than physically removed (`REMOVE_DUPLICATES=false`), in line with `GATK best practices` Van Der Auwera et al., 2013).

The process was run with lenient validation stringency (`VALIDATION_STRINGENCY=LENIENT`) to accommodate minor format inconsistencies. This step generated BAM files with marked duplicates and corresponding metrics files for each sample, which were used for downstream variant calling.

### 3.7. Variant Calling

For the identification of genetic variants (SNPs and indels) distinguishing the ‘Búlida Precoz’ mutant from its wild-type counterpart, the Genome `Analysis Toolkit (GATK, v4.6.2.0`) was employed (McKenna et al., 2010). Specifically, the `Mutect2` pipeline was selected (Benjamin et al., 2019) as it is a specialized somatic caller optimized for detecting the low-frequency mutations as expected in a bud sport (theoretically ∼16.7% VAF, see Introduction). This approach leverages a Bayesian statistical model designed to accurately call low-frequency alleles while filtering out germline variants and sequencing noise.

The workflow began with the creation of a Panel of Normals (PoN). The PoN was built by running `Mutect2` on all four wild-type samples in “tumor-only” mode and then using `CreateSomaticPanelOfNormals` to generate a single VCF file. The PoN serves as a database of recurrent technical artifacts, allowing `Mutect2` to filter these site-specific errors during variant calling.

The main variant calling was then performed for each mutant sample individually. Each mutant BAM file was provided as the “tumor” input, while a merged BAM file of the four wild-type samples was used as the “normal” control. By providing the previously generated PoN via the -pon flag, `Mutect2` called variants with high specificity, resulting in four raw VCF files, each containing potential somatic variants for one mutant replicate.

It is important to note that this initial variant calling stage does not, by itself, account for the significant differences in sequencing coverage observed between the samples (**Table S1**). The raw VCF files generated at this point are subsequently processed in a dedicated, multi-stage filtering workflow, as detailed in the following section. This step ensures that the number of reads supporting a variant call is evaluated relative to the mean coverage of that specific sample, providing a robust and equitable method for handling coverage variability.

### 3.8. Variant Filtering and Candidate Mutation Identification

Following the initial variant calling with `Mutect2`, a multi-stage filtering strategy was employed. This process was designed to systematically remove technical artifacts and to apply a stringent biological filter based on the experimental design, ultimately isolating a high-confidence list of candidate somatic mutations.

The workflow was performed for two strategies: one using `Mutect2’s` default filters alone, and a second that included an additional proportional read depth (DP) filter at the variant position to account for coverage variability between samples. First, `gatk FilterMutectCalls` was executed on the four raw `Mutect2` VCFs. This initial step applies `Mutect2’s` internal statistical models to tag common sequencing artifacts and unreliable calls, annotating each variant’s FILTER column. For the enhanced filtering strategy, a proportional DP filter of 15% of each sample’s mean coverage was then applied using `bcftools filter` (Danecek et al., 2021).

Finally, for each strategy, the filtered VCFs were merged and a strict biological filter was applied using `bcftools view`. The expression `N_PASS(GT=“alt”)==4` was used to select only those variants that were present in all four mutant replicates. Candidate mutations identified by these pipelines were then manually curated using the **Integrative Genomics Viewer (IGV)** (Robinson et al., 2011) to discard artifacts and confirm the variants.

### 3.9. Functional Annotation of Candidate Variants

To ascertain the biological significance of the final candidate variants identified by the `Mutect2` pipeline, a functional annotation step was performed using `SnpEff (v5.2)` (Cingolani et al., 2012). First, a custom `SnpEff` database was built for the project’s reference genome (`BUL_cur_guided.v1.0.fasta`) using its corresponding gene model information (`Bulida_annotationRNAseq.gff3`). Due to minor inconsistencies between the custom genome and its annotation file, the -`noCheckCds` and -`noCheckProtein` flags were utilized to ensure the successful creation of the database. Subsequently, the `snpEff ann` command was applied to the final candidate VCF file. This process annotated each variant with an ANN tag in the VCF’s INFO field, which contains critical information including the affected gene, the predicted functional effect (e.g., missense_variant -producing a different amino acid-, stop_gained -producing a stop codon-), and its putative impact (HIGH, MODERATE, LOW, or MODIFIER). Finally, all high-impact candidate mutations were manually curated using the Integrative Genomics Viewer (IGV) (Robinson et al., 2011) to discard potential artifacts and visually confirm the variants in the read alignments.

### 3.10. Comparative Analysis and Candidate Prioritization

A comparative analysis was designed to evaluate the two `Mutect2` filtering strategies (Default vs. Proportional DP) and to identify the highest-confidence candidate variants. The analysis was conducted using custom scripts in`Python` (v.3.0) (Van Rossum & Drake, 2009).

The final annotated data tables from each strategy were loaded using the `pandas` library (McKinney, 2010; The pandas development team, 2020). The primary comparison was performed at the gene level. The unique gene identifiers affected by variants in each of the two final callsets (Mutect2 Default, and Mutect2 15% DP) were extracted into `Python` set objects. The overlap between these sets was then systematically calculated and visualized using the `matplotlib-venn` library (Tretyakov, 2012/Tretyakov, 2025) to generate Venn diagrams. This provided a clear visual representation of the concordance between the two filtering strategies.

A final, prioritized list of candidate variants was then generated by selecting variants from the `Mutect2 15%` DP pipeline that had a predicted SnpEff impact of **HIGH** or **MODERATE**.

### 3.11. Functional Enrichment and Homology Analysis

The final phase of the analysis focused on elucidating the biological context of the prioritized candidate genes. This was achieved through a three-step workflow involving a homology search, Gene Ontology (GO) term annotation, and a statistical enrichment analysis.

First, a homology search was performed to infer the function of the candidate genes. The protein sequences of the candidates were used as queries in a local `BLASTp` (Altschul et al., 1990) search against a database created from the *Prunus armeniaca* reference proteome from UniProt. This step identified homologous proteins with known or predicted functions based on sequence similarity.

Next, to enrich these findings, the UniProt accession numbers from the top BLAST hits were used to retrieve all associated **Gene Ontology (GO) terms** for each candidate by querying the UniProt REST API.

Finally, a statistical over-representation test was performed to identify biological pathways or functions significantly affected by the mutations. This analysis used the `clusterProfiler` package (Yu et al., 2012) in R to compare the list of GO terms associated with the candidate genes against a background set representing all annotated genes in the genome.

## 4. RESULTS

This section details the findings from the bioinformatic analysis of whole-genome resequencing data derived from ‘Búlida’ wild-type (wt) and ‘Búlida Precoz’ mutant (mt) apricot samples. The primary aim is to identify candidate somatic mutations responsible for the early-flowering and low-chill phenotype of ‘Búlida Precoz’. The results presented will cover the initial data pre-processing and quality assessment stages, including read trimming, alignment outcomes, and genome coverage metrics. Subsequently, this section will elaborate on the outcomes of somatic variant calling using the `GATK Mutect2 pipeline`, followed by the filtering strategies applied to pinpoint high-confidence somatic variants differentiating the mutant from the wild-type. Data throughout this section are derived from the outputs of the various bioinformatic tools employed in the analytical pipeline, as detailed in Materials and Methods.

### 4.1. Sequencing Data Summary and Alignment Coverage

The Illumina paired-end sequencing of four biological replicates for ‘Búlida’ wild-type (wt1-wt4) and four for ‘Búlida Precoz’ mutant (mt1-mt4) yielded a substantial number of reads for analysis. Following adapter and quality trimming, the cleaned reads were aligned to the ‘Búlida’ reference genome. The total number of reads proceeding to alignment for each sample, along with the resulting mean depth of coverage across the ∼233 Mbp reference genome, are summarized in **Table S1**. These metrics were derived from the outputs of samtools flagstat (Danecek et al., 2021) and mosdepth (Pedersen & Quinlan, 2018).

### 4.2. PCR Duplicate Marking

Following alignment and BAM processing, duplicate reads were identified and marked using `Picard MarkDuplicates` (*Picard Tools - By Broad Institute*, s. f.) to mitigate potential biases in subsequent variant calling.

Prior to variant calling, it is essential to identify and account for duplicate reads that can arise from different stages of the sequencing workflow. These duplicates primarily fall into two categories: PCR duplicates and optical duplicates. PCR duplicates are generated during the PCR amplification step of library preparation, where a single DNA fragment is amplified multiple times. Optical duplicates, on the other hand, are artifacts of the Illumina sequencing process itself.`Picard MarkDuplicates` was used to identify both types, flagging them for downstream tools while retaining them in the BAM file. The key duplication metrics for each sample are summarized in **Table S2**.

**Key Observations from Duplication Metrics:**

- **Duplication Rates:** The percentage of duplication varied significantly between the two sequencing batches, a difference that correlates directly with their sequencing depth. Samples processed at the Genome Centre (wt1, wt2, mt1, mt2), which had a mean coverage of ∼85-103x, exhibited lower duplication rates, ranging from approximately 6.60% to 7.52%. In contrast, samples processed through outsourcing (wt3, wt4, mt3, mt4) showed substantially higher duplication rates, from 24.78% to 25.73%. This increase is consistent with their much deeper sequencing (∼202-248x), as a greater sequencing depth increases the probability of re-sequencing the same original DNA fragments. This effect may also be compounded by variations in the library preparation protocols between the two providers.
- **Optical Duplicates:** A fraction of the identified duplicates were classified as optical duplicates, distinct from PCR duplicates.
- **Estimated Library Size**: This metric, calculated by `Picard`, estimates the number of unique DNA fragments present in the original library before amplification. The values suggest reasonable library complexity for all samples, though outsourcing samples appear to have somewhat lower estimated library sizes relative to their higher duplication rates.

### 4.3. Sample Identity and Relationship Verification

To verify the genetic identity of all eight samples and rule out potential cross-contamination or sample swaps, a pairwise comparison was performed to quantify the genetic similarity between them. The methodology involved four main steps to transform the raw variant data into a visual representation.

First, a comprehensive pairwise comparison was conducted using the technically filtered, joint-called VCF file from the `HaplotypeCaller` workflow as input (*HaplotypeCaller*, 2025). The `bcftools isec` (Danecek et al., 2021) command was run on every possible pair of samples to generate raw counts of their shared and unique genetic variants. Second, these raw counts were converted into a standardized similarity score using the **Jaccard Index**. This metric was calculated for each pair by dividing the number of shared variants by the total number of unique variants present between them, yielding a score between 0 (no similarity) and 1 (identical).

Third, these Jaccard scores were used to construct a symmetrical 8×8 similarity matrix, with each sample represented as both a row and a column. Finally, this numerical matrix was visualized as a heatmap (**Figure S1**). The resulting heatmap visually confirms the integrity of the experimental design, showing two distinct high-similarity clusters corresponding to the mutant (mt1-mt4) and wild-type (wt1-wt4) replicate groups. Subtle sub-clustering is also visible (e.g., higher similarity between mt3/mt4 and wt3/wt4), a pattern consistent with the samples being processed in two different sequencing batches with different sequencing depth. Overall, this analysis robustly validates the sample identities, rules out potential swaps, and confirms the dataset is suitable for the subsequent differential variant analysis.

### 4.4. Variant Calling and Initial Filtering

The somatic variant calling pipeline, which compared each mutant sample against a pooled wild-type control using `GATK Mutect2`, produced two distinct sets of high-confidence somatic variants based on two parallel filtering strategies. The first strategy relied on `Mutect2’`s default filters, while the second, incorporated an additional filter based on read depth to account for coverage variability between samples.

To handle this variability and determine an optimal threshold, an iterative filtering experiment was conducted. This involved applying a filter that required the **total read depth (DP)** at a variant’s position to be at least a certain percentage of that sample’s mean genome-wide coverage. As shown in **Figure S2**, increasing the stringency of this proportional depth filter from **3% to 90%** leads to a clear and progressive reduction in the number of final candidates identified, from **123 candidates at 3%** to just **1 candidate at 90%**, discarding germline mutations. For all subsequent in-depth analyses, the **15% threshold with 112 variants** was chosen as it provides a robust balance between stringency and sensitivity, aligning with the theoretical expectation for detecting heterozygous somatic mutations. The sharp drop in candidates at the highest thresholds (only 8 at 70% and 1 at 90%) further supports this hypothesis, suggesting the variants are not homozygous or high-frequency artifacts.

Following the technical filtration, a biological filter was applied to isolate candidate mutations fixed within one sample but absent in the other. The analysis searched for two patterns: variants present in all four mutant replicates and absent from all four wild-type replicates (**4/4 vs. 0/4**), and the inverse pattern, variants present in all four wild-types but absent from all four mutants (**0/4 vs. 4/4**), which could represent Loss-of-Heterozygosity (LOH) events. The application of these workflows yielded significantly different results. The callset generated using only the default `Mutect2` filters contained **13**,**312** high-confidence somatic variants. In contrast, when the additional 15% proportional depth filter was applied, a more refined set containing **112** high-confidence somatic variants was obtained. This substantial reduction of over 99% underscores the critical impact of applying a coverage-aware depth filter for refining the final candidate list.

### 4.5. Functional Characterization of Mutect2 Candidates

A parallel analysis was conducted on the variants identified by the somatic caller, `GATK Mutect2` (Benjamin et al., 2019). To determine the most effective filtering strategy, two distinct approaches were compared: a standard workflow using only `GATK’s` default filters, and an enhanced workflow that incorporated an additional proportional read depth (DP) filter.

#### 4.5.1. Analysis of Candidates from the Default Filtering Strategy

The initial workflow, which relied solely on the filters applied by `FilterMutectCalls`, was first analyzed. After applying the strict biological filter (4/4 in mutants), a total of 13,312 candidate variants were identified. The functional annotation of this large set revealed that the vast majority were located in non-coding regions, classified with MODIFIER (**12**,**992 or ∼98%**), and 232 variants with HIGH or MODERATE impact (**Figure S3**). A breakdown of the specific functional effects further confirmed this, with `intergenic_region and upstream/downstream_gene_variant` being the most frequent categories (**Figure S4**). Although this initial set contained a subset of potentially interesting variants, the large total number strongly suggested a high proportion of false positives, likely arising from technical artifacts in regions of low coverage, making this dataset difficult to interpret and justifying the need for more stringent filtering.

#### 4.5.2. Analysis of Candidates from the Proportional Depth Filtering Strategy

To refine the candidate list, an additional filter requiring all mutant samples to have a read depth of at least 15% of their mean coverage at the variant site was applied. This more rigorous approach had a dramatic effect on the results. After applying the same strict 4/4 biological filter, the number of final candidates was reduced by over 99% to a final, manageable set of **112 high-confidence variants**. A detailed characterization of these variants from the **SnpEff summary report** revealed that the majority were Single Nucleotide Polymorphisms (SNPs), with a smaller fraction of insertions and deletions (indels) (**Figure 4**). The calculated transition-to-transversion ratio (Ts/Tv) for the SNPs was 1.9167, a value that indicates that there is a strong bias towards transitions when applying this filtering parameters (**Figure 5)**.

**Figure 4.**
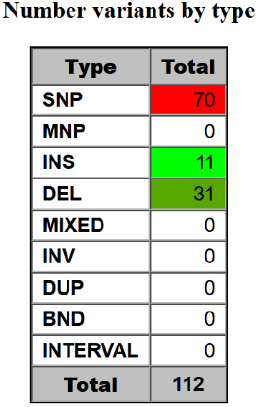
Variant Type Distribution for the Final Mutect2 Candidate Set. The chart shows the proportion of Single Nucleotide Polymorphisms (SNPs), Insertions (INS), Deletions (DEL), etc, among the 112 high-confidence candidates identified by the Mutect2 workflow.

**Figure 5.**
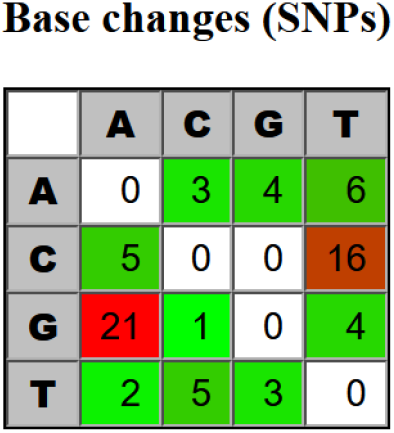
Nucleotide Substitution Profile of SNP Candidates. The grid shows the frequency of each of the 12 possible single nucleotide substitutions.

Functionally, the distribution of impacts for this refined set showed a clear and interpretable profile, with 1 HIGH impact, 3 MODERATE impact, and the remainder classified as LOW or MODIFIER. To validate these highest-priority candidates, a Variant Allele Frequency (VAF) analysis was performed. As shown in **Figure S5**, the high/moderate impact candidates display a consistent VAF clustering around the 0.5 threshold across all four mutant replicates. While this frequency is consistent with the theoretical expectation for a stable heterozygous somatic mutation, it is important to interpret this result with caution. Other biological scenarios could yield a similar allele frequency, for instance, a homozygous mutation resulting from a Loss-of-Heterozygosity (LOH) event within the majoritarian L2 cell layer could produce a VAF approaching this range. Furthermore, the identity and stability of meristematic layers are not always absolute, and cell migration between layers has been previously observed (Stewart & Dermen, 1975). Nevertheless, the strong consistency of the VAF across all four independent biological replicates provides compelling evidence that these variants represent somatic events stably maintained in the mutant lineage, rather than being technical artifacts.

This extensive filtering process ultimately distilled the candidate list to **four high-confidence variants** with HIGH or MODERATE functional impact. These variants, detailed in **Table 1**, represent the most robust somatic candidates identified by the `Mutect2` **pipeline**.

**Table 1.** High and Moderate Impact Variants Identified by the Mutect2 Pipeline.

|  |  |  |  |  |  |  |
| --- | --- | --- | --- | --- | --- | --- |
| Chromosome | Position | Reference Allele | Alternate Allele | Predicted Effect | Putative Impact | Affected Gene ID |
| BULcur1G | 9174501 | C | T | stop_gained | HIGH | Gene_9712 |
| BULcur4G | 9170462 | G | A | missense_variant | MODERATE | Gene_26305 |
| BULcur6G | 4620203 | G | T | missense_variant | MODERATE | Gene_20739 |
| BULcur7G | 15131066 | C | T | missense_variant | MODERATE | Gene_42548 |

The four high-confidence variants with a predicted HIGH or MODERATE functional impact were subjected to manual curation by visually inspecting the read alignments in the Integrative Genomics Viewer (IGV) (Robinson et al., 2011). As shown in **Figure S6**, the IGV snapshots for all four candidate loci reveal a clear and consistent pattern: the alternate allele is robustly supported by numerous reads in all four mutant replicates and is completely absent in all four wild-type replicates. This visual validation provides the ultimate confirmation that these are true, stable somatic mutations and not systematic artifacts of the bioinformatic pipeline.

#### 4.5.3. Identification of Potential Loss-of-Heterozygosity (LOH) Events

To identify variants potentially lost in the mutant lineage, an inverse biological filter was applied to the technically filtered, joint-called VCF file **generated by the** `GATK HaplotypeCaller` **workflow** (*HaplotypeCaller*, 2025) containing all eight samples. This analysis aimed to find variants present in all four wild-type replicates but absent from all four mutant replicates (a 0/4 vs. 4/4 pattern), which could represent Loss-of-Heterozygosity (LOH) events. Using `bcftools view` (Danecek et al., 2021), a filter `(‘N_PASS(GT[4-7]=“alt”) ==4 &∆ N_PASS(GT[0-3]=“ref”)==4’)` was applied to select variants where all wild-type samples carried the alternate allele and all mutant samples were homozygous for the reference allele. The resulting candidates were functionally annotated with `SnpEff`, and those with a predicted HIGH or MODERATE impact were prioritized. This workflow identified two high-confidence LOH candidates with moderate impact, which are detailed in **Table 2**.

**Table 2.**
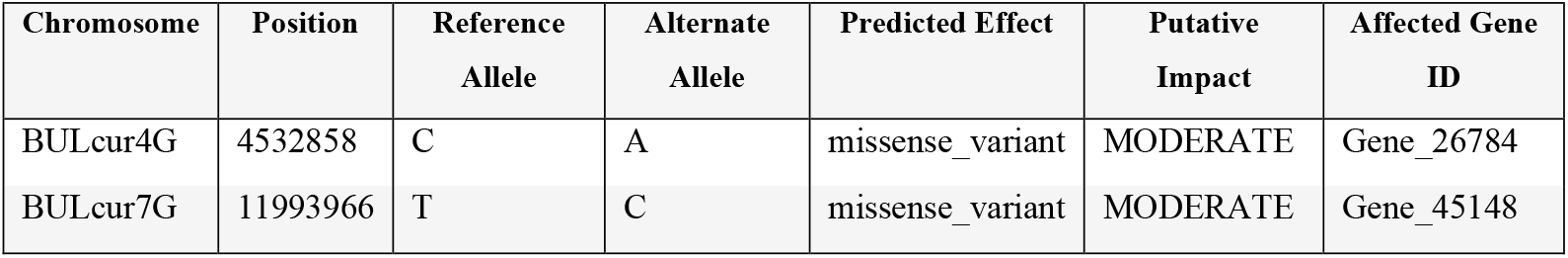
High/Moderate Impact LOH Candidate Variants.

### 4.6. Comparative Analysis and Identification of High-Confidence Genes

The final phase of the analysis was to identify the highest-confidence candidate genes. This was achieved by first comparing the two `Mutect2` filtering strategies (Benjamin et al., 2019) (Default vs. 15% DP) to select the most robust callset. Subsequently, the results from this enhanced `Mutect2` pipeline were cross-validated against an independent analysis using the `GATK HaplotypeCaller` (*HaplotypeCaller*, 2025) to provide an additional layer of algorithmic confirmation.

#### 4.6.1. Concordance Between Analytical Pipelines

Before the comparison, the list of unique genes affected by variants was extracted from each of the three analytical sets. Counting unique genes provides a more accurate functional overview than counting total variants. The results were as follows:

- The `HaplotypeCaller` **(15% DP)** pipeline identified 224 strict variants, which were located within **207 unique genes**.
- The `Mutect2` **(Default Filter)** pipeline identified a large set of strict variants affecting **2**,**832 unique genes**.
- The `Mutect2` **(15% DP Filter)** pipeline identified variants within **111 unique genes**.

The comparison between the two Mutect2 filtering strategies clearly shows that the proportional depth filter identifies a more refined subset of the genes found with the default filter (**Figure S7**). To further increase confidence in the Mutect2 (15% DP) callset, its results were compared against those from the `GATK HaplotypeCaller` pipeline. This cross-validation revealed a high degree of concordance, with a critical set of 102 genes being detected by both independent methodologies (**Figure 6**). A three-way comparison illustrates the complete relationship between all three callsets (**Figure S8**).

**Figure 6.**
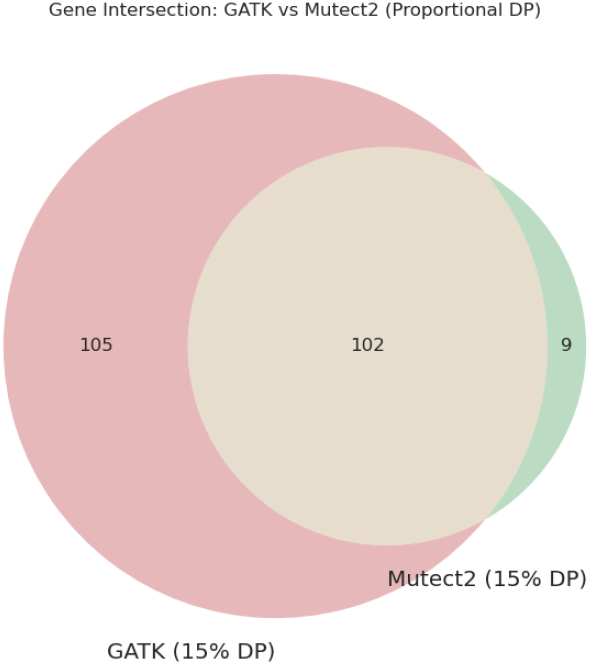
Comparison of Genes Identified by GATK HaplotypeCaller and the Mutect2 Pipeline. NcyThe diagram shows the overlap in affected genes between the final candidate sets from the GATK (207 genes) and Mutect2 15% DP (111 genes) pipelines.

#### 4.6.2. Final Prioritized List of Candidate Variants

The final list of prioritized candidates was defined as those variants with a predicted HIGH or MODERATE functional impact that were located in genes identified by the `Mutect2` (15% DP) pipeline and independently confirmed by the `GATK HaplotypeCaller` analysis. While `Mutect2` is the theoretically more precise tool for identifying somatic mutations, requiring this cross-algorithmic validation from a germline-focused caller provides the highest possible level of confidence in the results. This cross-validation resulted in a definitive list of four high-confidence genes.

The specific variants within these four genes are detailed in **Table 1**. These four variants, having been validated by independent algorithms and multiple layers of technical and biological filtering, represent the most promising candidates for harboring the causal mutation for the early flowering phenotype in ‘Búlida Precoz’.

### 4.7. Functional Annotation and Homology Analysis of Candidate Genes

To infer the potential biological function of the final high-confidence candidate genes, a homology-based annotation was performed. The protein sequences of the candidates were used as queries in a local **BLASTp** (Altschul et al., 1990) search against a local database created from the *Prunus armeniaca* reference proteome (UniProt organism ID: 36596). For each query, the single best hit was selected based on the highest bitscore to ensure the most biologically relevant match.

The results of this homology analysis are summarized in **Table 3**. All candidate genes showed high-quality alignments to known or predicted proteins in *Prunus armeniaca*. The alignments demonstrated high percentage identity and extremely low `E-values`, indicating that the sequence similarity is statistically significant and not a result of chance. This high degree of homology allows for a confident inference of function.

**Table 3.** Functional Annotation of Top Candidate Genes via BLASTp Homology.

| Candidate Gene ID | Predicted Protein Description | Percent Identity (%) | Bitscore | E-value |
| --- | --- | --- | --- | --- |
| Gene_9712 | Peroxiredoxin | 100.00 | 150.0 | 6.75e-48 |
| Gene_20739 | DNA-directed RNA polymerase I subunit rpa49 | 100.00 | 145.0 | 1.28e-43 |
| Gene_26305 | Methyltransferase domain-containing protein | 100.00 | 155.0 | 6.82e-47 |
| Gene_42548 | RRM domain-containing protein | 100.00 | 140.0 | 6.76e-41 |

Notably, the candidates include proteins with functions related to fundamental cellular processes. For example, **Gene_9712** was identified as a **Peroxiredoxin**, a key enzyme family involved in responding to oxidative stress. **Gene_20739** was identified as a subunit of **DNA-directed RNA polymerase I**, central to the process of transcription. **Gene_42548** corresponds to an **RRM domain-containing protein**, a family typically associated with RNA binding and post-transcriptional regulation. Finally, **Gene_26305** was annotated as a **Methyltransferase domain-containing protein**, a broad class of enzymes involved in many regulatory pathways. The identification of these conserved proteins suggests that the identified somatic mutations may be impacting core cellular functions, which could have cascading effects on the plant’s developmental timing.

### 4.8. Gene Ontology (GO) Enrichment Analysis

To understand the collective biological role of the final candidate genes, a Gene Ontology (GO) analysis was performed to identify biological functions or pathways that were disproportionately affected by the identified mutations. An initial descriptive analysis calculated the frequency of all GO terms associated with the candidate genes (**Table S3**), revealing a prevalence of terms related to fundamental molecular activities such as “RNA binding” (GO:0003723) and cellular locations like “nucleus” (GO:0005634). The distribution of the most frequent terms is visualized in **Figure S9**.

To determine if these functions were statistically significant, a GO enrichment analysis was performed using the R package `clusterProfiler` (Yu et al., 2012). This test compares the prevalence of GO terms in the candidate list against a genomic background to find over-represented terms (FDR < 0.05). The analysis yielded no significantly enriched terms for the Molecular Function (MF) ontology, a likely consequence of the small candidate set size.

However, the test for Cellular Component (CC) yielded two significant terms related to the spliceosome: *precatalytic spliceosome* (GO:0005681; adj. p ≈ 0.027) and *catalytic step 2 spliceosome* (GO:0005680; adj. p ≈ 0.030). Both of these terms were driven by the candidate gene **Gene_42548 (RRM domain-containing protein)** (**Figure S10**). For Biological Process (BP), seven terms were found to be significantly enriched (**Figure S11**). The results reinforce the findings from the descriptive analysis, with the most robust signal being the involvement of **Gene_42548** in RNA-processing, including *regulation of mRNA splicing, via spliceosome* (GO:0048024; adj. p ≈ 0.042). Additionally, **Gene_20739 (DNA-directed RNA polymerase I subunit)** was linked to *DNA-templated transcription* (GO:0006351; adj. p ≈ 0.042), while **Gene_9712 (Peroxiredoxin)** was associated with *cell redox homeostasis* (GO:0045454; adj. p ≈ 0.042).

These statistical results strongly suggest that the identified mutations have preferentially targeted genes involved in core informational and regulatory pathways, particularly RNA splicing and transcription, as well as cellular stress response. This provides a compelling hypothesis for the molecular basis of the ‘Búlida Precoz’ phenotype.

## 5. DISCUSSION

This study aimed to identify candidate somatic mutations responsible for the low-chill requirement and early-flowering phenotype of the ‘Búlida Precoz’ apricot mutant. The preceding sections have detailed the methodological pipeline applied, from raw sequencing data pre-processing through to a somatic variant calling workflow using `GATK Mutect2` (Benjamin et al., 2019) and a multi-stage filtering strategy aimed at isolating high-confidence candidate mutations. This section will now interpret the comprehensive findings of this study, relating them to the established biological context and the initial objectives. The significance of the identified candidate somatic mutations, their potential impact on understanding flowering time regulation, and their practical applications for apricot breeding will be explored. Finally, methodological choices, inherent limitations of the study, and pertinent future research directions will be addressed.

### 5.1. Data Quality and Pre-processing Outcomes

The initial stages of the bioinformatic pipeline, encompassing raw read quality control, trimming, alignment, and PCR duplicate marking, are foundational for the reliability of subsequent variant discovery. The quality assessment of the raw Illumina paired-end sequencing data yielded high mapping rates for all samples, consistently exceeding 98.9%. This high percentage indicates minimal contamination and highlights the importance of using a close reference to the studied focal genome. Furthermore, the percentage of reads that mapped as “properly paired” was consistently high across all samples (92% to 97%), reflecting the integrity of the sequencing libraries and the structural consistency between the samples and the reference assembly.

Coverage analysis performed with mosdepth (Pedersen & Quinlan, 2018) revealed excellent mean depth across all samples. Two distinct groups of samples were observed based on sequencing depth. The first group (wt1, wt2, mt1, mt2) achieved mean coverages ranging from approximately 86x to 103x, while the second group (wt3, wt4, mt3, mt4) exhibited deeper coverage, from approximately 202x to 248x. Both coverage levels are well above the >50-60x depth often recommended for robust somatic variant detection, providing a strong foundation for identifying low-frequency mutations. An interesting artifact observed was the consistently higher mean coverage for the unplaced scaffold contig BULcur0G. This phenomenon is likely due to the ambiguous mapping of repetitive sequences from the main chromosomes to this artificial chromosome, artificially inflating its calculated coverage depth.

For this study, the workflow was transitioned to a post-alignment strategy using `Picard MarkDuplicates` (*Picard Tools - By Broad Institute*, s. f.), in line with GATK Best Practices for WGS (Van Der Auwera et al., 2013). This approach is more robust for WGS data as it identifies duplicates by their mapping coordinates, correctly handling reads that are identical in origin but contain minor sequencing errors. This adaptation, while a deviation from the reference workflow, ensures a more accurate handling of PCR and optical duplicates, thereby improving the reliability of the final somatic variant calls.

The level of PCR duplication, marked with `Picard MarkDuplicates`, correlated with sequencing depth. The samples with ∼86-103x coverage showed lower duplication rates (approximately 6.6% to 7.5%) compared to the samples with >200x coverage (approximately 24.8% to 25.7%). These differences likely reflect variations in library preparation protocols or initial DNA input amounts between the two sequencing batches. Despite the higher duplication in the deeper-sequenced set, the estimated library sizes for all samples suggest sufficient unique molecular complexity for downstream analyses. Overall, the pre-processing and alignment stages yielded high-quality data suitable for the subsequent steps of somatic mutation discovery.

### 5.2. Annotation

For the functional annotation of variants, `SnpEff` was chosen over other tools like `GATK’s Funcotator`. While `Funcotator` is the recommended annotator within the `GATK` framework, particularly for human somatic studies, `SnpEff` offers significant advantages for projects involving non-model organisms with custom genomes. The primary reason for its selection was its straightforward and well-documented workflow for building a custom annotation database from a reference FASTA and a GFF3 file, a necessary prerequisite for this project. This flexibility ensured an accurate and reliable annotation based specifically on the ‘Búlida’ reference genome.

#### 5.3. GATK Mutect2 for Somatic Variant Calling in Bud Sports

The identification of somatic mutations in bud sports was conducted using `GATK Mutect2` (Benjamin et al., 2019), which is specifically designed for identifying variants in underrepresented lineages in cancer genetics, an anologous model to the meristematic layers in budsports. Its primary advantage is its ability to directly compare a “tumor” sample (the mutant) against a matched “normal” sample (the wild-type pool), which is crucial for distinguishing true somatic events from background germline polymorphisms. Furthermore, Mutect2 models allele fractions rather than assuming a fixed ploidy, making it highly sensitive to the low-frequency variants characteristic of somatic mosaicism in different meristematic layers (e.g., L1, L2, L3). The use of a Panel of `Normals (PoN)` constructed from the wild-type replicates further enhanced specificity by allowing the filtering of systematic technical artifacts.

A critical step in the Mutect2 workflow was the application of the filtering strategy. While `Mutect2’s` default filters yielded a large set of 13,312 candidate genes, the addition of a 15% proportional depth filter proved essential. This extra step refined the results by over 99%, producing a high-confidence list of variants affecting just **111** unique genes, effectively removing likely false positives from low-coverage regions.

To provide the highest level of confidence in these final candidates, an independent, cross-algorithmic validation was performed. The 111 genes identified by the Mutect2 pipeline were compared against a candidate set generated by the germline caller `GATK HaplotypeCaller` (*HaplotypeCaller*, 2025). A remarkable degree of concordance was observed, with the vast majority of the genes identified by Mutect2 also being detected by the `HaplotypeCaller` pipeline (**Figure 6**). This high level of agreement, where a less specialized tool confirms the findings of the purpose-built somatic caller, lends exceptional robustness to the results and strongly suggests that the identified candidate genes represent true biological signals. The genes present in this intersection are thus considered the definitive candidates for harboring the causal mutation.

A preliminary analysis of the variant callsets revealed a significant challenge: the mean sequencing coverage varied considerably across the eight samples, ranging from approximately 86x to over 248x. This heterogeneity posed a risk, as applying a single, fixed read depth (DP) filter across all samples could disproportionately penalize variants in lower-coverage samples. To address this, a coverage-aware, proportional filtering strategy was designed and implemented.

The core of this workflow was an iterative experiment that automated the testing of several distinct filtering stringencies, based on a percentage of each sample’s mean coverage (from 3% to 50%). For each percentage level, a specific minimum DP threshold was calculated for every one of the eight samples. Subsequently, a chained pipeline of `bcftools filter` (Danecek et al., 2021) commands was used to apply these proportional DP filters in addition to standard quality filters.

To evaluate the impact of this filtering, the strict biological filter (4/4 in mutants and 0/4 in wild-types) was applied to each of the technically filtered VCF files. The results clearly demonstrated a direct correlation between filtering stringency and the final number of high-confidence candidates. This empirical approach not only allowed for the selection of the 15% threshold as the most robust choice for the final analysis but also served as a crucial quality control step. The reduction in candidate numbers observed in the `Mutect2` pipeline confirmed that this proportional filtering was essential for effectively removing low-coverage artifacts and isolating a reliable set of somatic variants for downstream biological interpretation.

### 5.4. Mutational Signatures

A detailed characterization of the variants identified by the `Mutect2` pipeline (15% proportional DP filter) revealed a calculated transition-to-transversion (Ts/Tv) ratio for the identified SNPs of **1.9167**. This value, indicating a strong bias towards transitions, serves as an important internal quality control metric and aligns with known mutational patterns.

This prevalence of transitions is highly consistent with patterns of somatic mutation reported in other *Prunus* species. For instance, in apricot (*Prunus armeniaca*), a strong bias towards C to T and G to A transitions was observed in somatic point mutations (SNPs) found in the L1 (epidermis) and L2 (mesocarp) layers of fruits (Goel et al., 2024). Similarly, in apple (*Malus domestica* cv. McIntosh RubyMac), *de novo* mutations (gain of heterozygosity or GOH) showed a notable enrichment of GC to AT transitions, a spectrum consistent with inherited germline mutations (Sun et al., 2024). The identification of both SNP and indel mutations also aligns with variants analyzed in these studies, which include indels and loss of heterozygosity (LOH) events or gene conversions as types of somatic changes (Goel et al., 2024; Sun et al., 2024).

Furthermore, the observation that a majority of somatic mutations are layer-specific in apricot (94%) and that a significant proportion (approximately 20%) is also layer-specific in apple, highlights the critical importance of considering the meristematic organization of plants when analyzing genomic mosaicism. This is particularly relevant as, for clonal crops, the representativeness of cell layers and the sequencing depth (often requiring >50x-60x coverage for low allelic frequency somatic mutations) are crucial for distinguishing true somatic events from sequencing errors. The prevalence of somatic mutations in L1 (epidermis) over L2 (mesocarp) in apricot and L3 over L2 in apple further underscores the distinct mutational dynamics within different meristematic layers (Goel et al., 2024; Sun et al., 2024). This is of great importance as only L2 mutations will be transmitted to the next generation, as gametes are produced from L2 tissue.

### 5.5. Biological Interpretation of High-Confidence Candidate Genes

The final objective of this study was to identify the genetic basis of the early flowering phenotype in ‘Búlida Precoz’. The bioinformatic workflow culminated in a short, robust list of candidate genes. The interpretation of their biological functions, supported by the Gene Ontology (GO) enrichment analysis, allows for a molecular model to be proposed to explain this agronomically valuable phenotype. The functional enrichment analysis revealed two primary biological signals: a potential alteration in cellular redox homeostasis and a disruption of the machinery for RNA processing and gene expression.

The strongest signal related to cellular stress comes from **Gene_9712**, which harbors a stop_gained mutation. This type of mutation is particularly severe, as it leads to the production of a truncated and likely non-functional protein. Homology-based annotation identifies this gene as a Peroxiredoxin, a key enzyme in the detoxification of reactive oxygen species (ROS).

The link between redox state and dormancy release is well-documented and provides a hypothesis for the ‘Búlida Precoz’ phenotype. Studies on bud dormancy in peach have shown that the transition from endodormancy to ecodormancy is marked by a significant accumulation of hydrogen peroxide (H_₂_O_₂_), which acts as a key signaling molecule for bud break. This oxidative burst is tightly regulated by the activity of antioxidant enzymes, including peroxidases. It is therefore highly plausible that the loss-of-function mutation in the Peroxiredoxin identified in this study disrupts this delicate redox balance. An inability to properly regulate H_₂_O_₂_ levels could lead to its premature accumulation, effectively sending an “early wake-up call” to the buds and triggering an earlier exit from dormancy, which would directly result in the observed early flowering phenotype (Considine & Foyer, 2024; Hernandez et al., 2021).

The second primary biological signal identified by the functional enrichment analysis points to a significant dysregulation of gene expression processes, providing a compelling hypothesis for the early-flowering phenotype. **Gene_42548**, annotated as an RRM domain-containing protein, emerges as a central candidate. The enrichment analysis statistically implicated this gene in regulation of mRNA splicing, via spliceosome (GO:0048024) and in localization to the spliceosome complex itself (GO:0005681, GO:0005680). This finding is particularly significant, as the link between alternative splicing and the control of flowering time is well-established in plants. For instance, a study in *Arabidopsis thaliana* demonstrated that U2AF65, an RNA-binding protein containing RRM domains and a key component of the spliceosome, directly regulates flowering time by controlling the alternative splicing of key floral repressor genes like *FLOWERING LOCUS C* (*FLC*) (Park et al., 2019). The discovery of a mutation in a core splicing factor like Gene_42548 in ‘Búlida Precoz’ suggests a highly plausible mechanism where altered splicing of such developmental regulators could disrupt their normal function, thereby promoting an early transition to flowering. This is further supported by studies in closely related species: 1) *Prunus avium*, in which it has been identified alternative splicing in genotypes with different chilling requirements (Martínez-Romera et al., 2025). 2) *Prunus persica*, in which it has been identified genes involved in “RNA processing” and “transcription” as being differentially expressed during the release from bud dormancy (Leida et al., 2010).

This hypothesis is further strengthened by the identification of a mutation in **Gene_20739**, annotated as a DNA-directed RNA polymerase I subunit and linked to the GO term DNA-templated transcription (GO:0006351). This suggests that the impact on gene expression may not be limited to post-transcriptional processing but could also involve the act of transcription itself. While RNA Polymerase I is primarily responsible for synthesizing ribosomal RNA, its activity is fundamentally linked to the cell’s overall metabolic state and capacity for growth. The regulation of ribosome biogenesis is known to be integrated with developmental pathways, as the cell must be prepared for the protein synthesis demands of new growth (Chen et al., 2012). Therefore, an alteration in this core transcriptional machinery could influence the timing of major developmental transitions like bud break and flowering.

Taken together, these results suggest a model in which the ‘Búlida Precoz’ phenotype is not caused by a mutation in a single, canonical “flowering gene,” but rather by a disruption in fundamental cellular processes. The dysregulation of redox homeostasis and, more prominently, of the transcription and splicing machinery, could be causing a global reprogramming of gene expression that ultimately manifests as an acceleration of development and early flowering.

### 5.6. Significance for Apricot Adaptation and Breeding

The identification of high-confidence candidate genes responsible for the low-chill, early-flowering phenotype in the ‘Búlida Precoz’ has significant implications for both our fundamental understanding of plant development and the practical application of apricot breeding programs. As traditional temperate fruit cultivation is increasingly threatened by climate change and insufficient winter chilling, understanding and utilizing new genetic resources for adaptation has become a critical priority. This study serves as a powerful case study, demonstrating how a single (or several) somatic mutation(s) can lead to an agronomically valuable trait. The findings suggest that the mechanism for adapting to lower chilling requirements may not lie within the canonical flowering-time pathways alone but can also arise from alterations to fundamental cellular processes such as RNA processing and cellular redox homeostasis. The identification of candidate genes like a putative RNA-binding protein (Gene_42548) and a Peroxiredoxin (Gene_9712) opens new avenues for research into the complex regulatory networks that govern dormancy and floral transition in response to environmental cues.

From a practical standpoint, the most significant outcome of this research is the potential for marker-assisted breeding. The development of new apricot cultivars through traditional breeding is a slow, resource-intensive process, often requiring a decade or more to evaluate new selections. The specific, high-confidence variants identified in this study, particularly the stop_gained mutation, can be converted into robust molecular markers following their validation in a ‘Búlida Precoz’-derived progeny. These markers would allow breeders to screen thousands of seedlings at a very early stage, rapidly identifying individuals that carry the desirable low-chill trait without waiting years for the trees to overcome juvenility. This would dramatically accelerate the breeding cycle, enabling a more rapid development of new, climate-resilient apricot varieties adapted to warmer winter conditions. Ultimately, this work provides not only a list of candidate genes but also the foundational tools for future breeding efforts aimed at ensuring the sustainability of apricot cultivation.

### 5.7. Future Perspectives

While this study successfully identified a robust set of high-confidence candidate variants, it is essential to acknowledge its limitations and the future work required for validation. The findings are entirely computational, and the causal link between the identified variants and the early-flowering phenotype remains a hypothesis until experimentally confirmed. Furthermore, the analysis is contingent on the quality of the reference genome and its annotation. A key consideration is the limitation imposed by sequencing depth; while robust, the ∼100x average coverage may have been insufficient to confidently detect mutations exclusive to underrepresented meristematic layers like the L1, which are reported to be frequent in apricot (Goel et al., 2024). Future work must therefore focus on the experimental validation of the top candidate variants, for instance by confirming their presence via Sanger sequencing and assessing their functional impact on gene expression using RT-qPCR. These subsequent steps will be crucial to move from the computational predictions of this study to definitively uncovering the molecular mechanism behind this agronomically valuable trait.

## 6. CONCLUSIONS

The bioinformatic analysis conducted in this study successfully identified the most probable genetic determinants of the early flowering phenotype in the ‘Búlida Precoz’ apricot. A rigorous, multi-stage filtering workflow effectively reduced over 13,000 initial raw variants to a final, high-confidence list of four candidate genes containing somatic mutations. Within this final list, the identification of a **stop_gained (high-impact) mutation in the gene Gene_9712**, annotated as a **Peroxiredoxin**, stands out as the most promising candidate to explain the phenotype, given its functional implication in cellular stress response. Furthermore, the functional enrichment analysis of all candidate genes suggests that the early-flowering trait may not be caused by a mutation in a canonical flowering gene, but rather by a potential dysregulation of fundamental cellular processes. Primarily, these processes are **redox homeostasis** (linked to the Peroxiredoxin) and **RNA processing and splicing** (linked to other high-confidence candidates like Gene_42548), opening new avenues for understanding developmental regulation in temperate fruit trees.

## 8. ACKNOWLEDGEMENTS

The authors would like to thank José Egea for kindly providing plant material, Birgit Walkemeier for DNA extraction and sample preparation, and Bruno Huettel for the assistance in the sequencing at the Max Planck Genome Centre. The help in the conceptualization of the study and preliminary analysis of the data by Hequan Sun (Xi’an Jiaotong University), Manish Goel, and Korbinian Schneeberger (Ludwig-Maximilians-Universität München) is gratefully acknowledged. The authors would also like to thank Bruno Contreras for helpful comments on the manuscript.

## 9. CODE AND DATA AVAILABILITY

To ensure complete transparency and reproducibility, all command-line scripts developed and used for this study are publicly available in the project’s GitHub repository: https://github.com/DanielGP121/Bulida-Precoz-Preprint. All supplementary tables and figures are provided in the Supplementary Material document, also available in this repository.

Computational analyses were performed on a Linux server with 10 CPU cores and 64 GB of RAM at the Estación Experimental del Aula Dei (EEAD-CSIC). Software installation and script execution were performed within a dedicated Conda environment, ensuring a consistent and reproducible software environment.

## REFERENCES

Altschul, S. F., Gish, W., Miller, W., Myers, E. W., & Lipman, D. J. (1990). Basic Local Alignment Search Tool. Babraham Bioinformatics—FastQC A Quality Control tool for High Throughput Sequence Data. (s. f.). Recuperado 24 de julio de 2025, de https://www.bioinformatics.babraham.ac.uk/projects/fastqc/

Benjamin, D., Sato, T., Cibulskis, K., Getz, G., Stewart, C., & Lichtenstein, L. (2019). Calling Somatic SNVs and Indels with Mutect2. 10.1101/861054

Campoy, J. A., Sun, H., Goel, M., Jiao, W.-B., Folz-Donahue, K., Wang, N., Rubio, M., Liu, C., Kukat, C., Ruiz, D., Huettel, B., & Schneeberger, K. (2020). Gamete binning: Chromosome-level and haplotype-resolved genome assembly enabled by high-throughput single-cell sequencing of gamete genomes. Genome Biology, 21(1), 306. 10.1186/s13059-020-02235-5

Chen, S.-Y., Ho, K.-J., Hsieh, Y.-J., Wang, L.-T., & Mau, J.-L. (2012). Contents of lovastatin, γ-aminobutyric acid and ergothioneine in mushroom fruiting bodies and mycelia. LWT, 47(2), 274–278. 10.1016/j.lwt.2012.01.019

Cingolani, P., Platts, A., Wang, L. L., Coon, M., Nguyen, T., Wang, L., Land, S. J., Lu, X., & Ruden, D. M. (2012). A program for annotating and predicting the effects of single nucleotide polymorphisms, SnpEff: SNPs in the genome of Drosophila melanogaster strain w^1118^; iso-2; iso-3. Fly, 6(2), 80–92. 10.4161/fly.19695

Considine, M. J., & Foyer, C. H. (2024). Redox regulation of meristem quiescence: Outside/in. Journal of Experimental Botany, 75(19), 6037–6046. 10.1093/jxb/erae161

Danecek, P., Bonfield, J. K., Liddle, J., Marshall, J., Ohan, V., Pollard, M. O., Whitwham, A., Keane, T., McCarthy, S. A., Davies, R. M., & Li, H. (2021). Twelve years of SAMtools and BCFtools. GigaScience, 10(2). 10.1093/gigascience/giab008

Goel, M., Campoy, J. A., Krause, K., Baus, L. C., Sahu, A., Sun, H., Walkemeier, B., Marek, M., Beaudry, R., Ruiz, D., Huettel, B., & Schneeberger, K. (2024). The vast majority of somatic mutations in plants are layer-specific. Genome Biology, 25(1). 10.1186/s13059-024-03337-0

HaplotypeCaller. (2025, julio 1). GATK. https://gatk.broadinstitute.org/hc/en-us/articles/360037225632-HaplotypeCaller

Hernandez, J. A., Díaz-Vivancos, P., Martínez-Sánchez, G., Alburquerque, N., Martínez, D., Barba-Espín, G., Acosta-Motos, J. R., Carrera, E., & García-Bruntón, J. (2021). Physiological and biochemical characterization of bud dormancy: Evolution of carbohydrate and antioxidant metabolisms and hormonal profile in a low chill peach variety. Scientia Horticulturae, 281, 109957. 10.1016/j.scienta.2021.109957

Jiang, H., Lei, R., Ding, S.-W., & Zhu, S. (2014). Skewer: A fast and accurate adapter trimmer for next-generation sequencing paired-end reads. BMC Bioinformatics, 15(1). 10.1186/1471-2105-15-182

Leida, C., Terol, J., Marti, G., Agusti, M., Llacer, G., Badenes, M. L., & Rios, G. (2010). Identification of genes associated with bud dormancy release in Prunus persica by suppression subtractive hybridization. Tree Physiology, 30(5), 655–666. 10.1093/treephys/tpq008

Li, H., & Durbin, R. (2009). Fast and accurate short read alignment with Burrows–Wheeler transform. Bioinformatics, 25(14), 1754–1760. 10.1093/bioinformatics/btp324

Martínez-Romera, N., Wünsch, A., & Hedhly, A. (2025, mayo 6). Sweet Cherry Dormancy: Characterization of PavDAMs Alternative Splicing in Genotypes with Different Chilling Requirements.

McKenna, A., Hanna, M., Banks, E., Sivachenko, A., Cibulskis, K., Kernytsky, A., Garimella, K., Altshuler, D., Gabriel, S., Daly, M., & DePristo, M. A. (2010). The Genome Analysis Toolkit: A MapReduce framework for analyzing next-generation DNA sequencing data. Genome Research, 20(9), 1297–1303. 10.1101/gr.107524.110

McKinney, W. (2010). Data Structures for Statistical Computing in Python. 56–61. 10.25080/Majora-92bf1922-00a

Park, H.-Y., Lee, H. T., Lee, J. H., & Kim, J.-K. (2019). Arabidopsis U2AF65 Regulates Flowering Time and the Growth of Pollen Tubes. Frontiers in Plant Science, 10. 10.3389/fpls.2019.00569

Pedersen, B. S., & Quinlan, A. R. (2018). Mosdepth: Quick coverage calculation for genomes and exomes. Bioinformatics, 34(5), 867–868. 10.1093/bioinformatics/btx699

Picard Tools—By Broad Institute. (s. f.). Recuperado 30 de mayo de 2025, de https://broadinstitute.github.io/picard/

Robinson, J. T., Thorvaldsdóttir, H., Winckler, W., Guttman, M., Lander, E. S., Getz, G., & Mesirov, J. P. (2011). Integrative genomics viewer. Nature Biotechnology, 29(1), 24–26. 10.1038/nbt.1754

Ruiz, D., García-Gómez, B. E., Egea, J., Molina, A., Martínez-Gómez, P., & Campoy, J. A. (2019). Phenotypical characterization and molecular fingerprinting of natural early-flowering mutants in apricot (Prunus armeniaca L.) and Japanese plum (P. salicina Lindl.). Scientia Horticulturae, 254, 187–192. 10.1016/j.scienta.2019.05.002

Somatic calling is NOT simply a difference between two callsets. (2024, junio 25). GATK. https://gatk.broadinstitute.org/hc/en-us/articles/360035890491-Somatic-calling-is-NOT-simply-a-difference-between-two-callsets

Stewart, R. N., & Dermen, H. (1975). FLEXIBILITY IN ONTOGENY AS SHOWN BY THE CONTRIBUTION OF THE SHOOT APICAL LAYERS TO LEAVES OF PERICLINAL CHIMERAS. American Journal of Botany, 62(9), 935–947. 10.1002/j.1537-2197.1975.tb14134.x

Sun, H., Abeli, P., Campoy, J. A., Rütjes, T., Krause, K., Jiao, W.-B., Beaudry, R., & Schneeberger, K. (2024). The identification and analysis of meristematic mutations within the apple tree that developed the RubyMac sport mutation. BMC Plant Biology, 24(1). 10.1186/s12870-024-05628-x

The pandas development team. (2020). pandas-dev/pandas: Pandas (Versión latest) [Software]. Zenodo. 10.5281/zenodo.3509134

Tretyakov, K. (2025). Konstantint/matplotlib-venn [Jupyter Notebook]. https://github.com/konstantint/matplotlib-venn (Obra original publicada en 2012)

Van Der Auwera, G. A., Carneiro, M. O., Hartl, C., Poplin, R., Del Angel, G., Levy‐Moonshine, A., Jordan, T., Shakir, K., Roazen, D., Thibault, J., Banks, E., Garimella, K. V., Altshuler, D., Gabriel, S., & DePristo, M. A. (2013). From FastQ Data to High‐Confidence Variant Calls: The Genome Analysis Toolkit Best Practices Pipeline. Current Protocols in Bioinformatics, 43(1). 10.1002/0471250953.bi1110s43

Van Rossum, G., & Drake, F. L. (2009). Python 3 Reference Manual. CreateSpace.

Yu, G., Wang, L.-G., Han, Y., & He, Q.-Y. (2012). clusterProfiler: An R Package for Comparing Biological Themes Among Gene Clusters. OMICS: A Journal of Integrative Biology, 16(5), 284–287. 10.1089/omi.2011.0118

