## Supplementary Material for "A stop-gained mutation in a Peroxiredoxin gene may underlie the low-chill phenotype of the apricot spontaneous mutant ‘Búlida Precoz’"

### Supplementary Tables

**Supplementary Table S1. Summary of Sequencing Data Used for Alignment and Resulting Genome Coverage.**

| Sample Name | Reads for Alignment (Post-Trimming) | Mean Depth of Coverage (x) |
|---|---|---|
| **wt1** | 179,550,177 | 103.07 |
| **wt2** | 179,316,392 | 102.53 |
| **wt3** | 356,835,396 | 210.11 |
| **wt4** | 339,235,619 | 202.32 |
| **mt1** | 180,564,548 | 103.02 |
| **mt2** | 151,058,488 | 85.66 |
| **mt3** | 354,588,337 | 213.94 |
| **mt4** | 413,499,653 | 247.94 |

**Supplementary Table S2. Duplication Metrics from Picard MarkDuplicates**

| Sample Name | Read Pairs Examined | Read Pair Duplicates | Read Pair Optical Duplicates | Percent Duplication (%) | Estimated Library Size |
|---|---|---|---|---|---|
| **wt1** | 88,103,136 | 5,787,039 | 454,979 | 6.72 | 690,855,889 |
| **wt2** | 87,443,228 | 5,739,281 | 448,684 | 6.76 | 685,933,940 |
| **wt3** | 176,491,955 | 43,927,291 | 2,918,546 | 24.97 | 306,854,928 |
| **wt4** | 167,361,099 | 41,290,705 | 2,928,143 | 24.78 | 295,146,062 |
| **mt1** | 88,020,817 | 6,470,363 | 666,428 | 7.52 | 627,928,134 |
| **mt2** | 73,436,523 | 4,704,557 | 485,302 | 6.60 | 606,106,918 |
| **mt3** | 175,182,685 | 44,714,326 | 3,170,265 | 25.63 | 296,105,958 |
| **mt4** | 204,295,644 | 52,362,696 | 3,637,542 | 25.73 | 343,158,453 |

**Supplementary Table S3. Top 10 Most Frequent GO Terms Associated with Candidate**

| GO Term ID | Frequency | Predicted GO Term Name |
|---|---|---|
| GO:0003723 | 3 | RNA binding |
| GO:0005634 | 3 | nucleus |
| GO:0006397 | 3 | mRNA processing |
| GO:0008380 | 3 | RNA splicing |
| GO:0045454 | 2 | cell redox homeostasis |
| GO:0005730 | 2 | nucleolus |
| GO:0003677 | 2 | DNA binding |
| GO:0000428 | 2 | pre-rRNA processing |
| GO:0006351 | 2 | transcription, DNA-templated |
| GO:0005829 | 1 | cytosol |

### Supplementary Figures

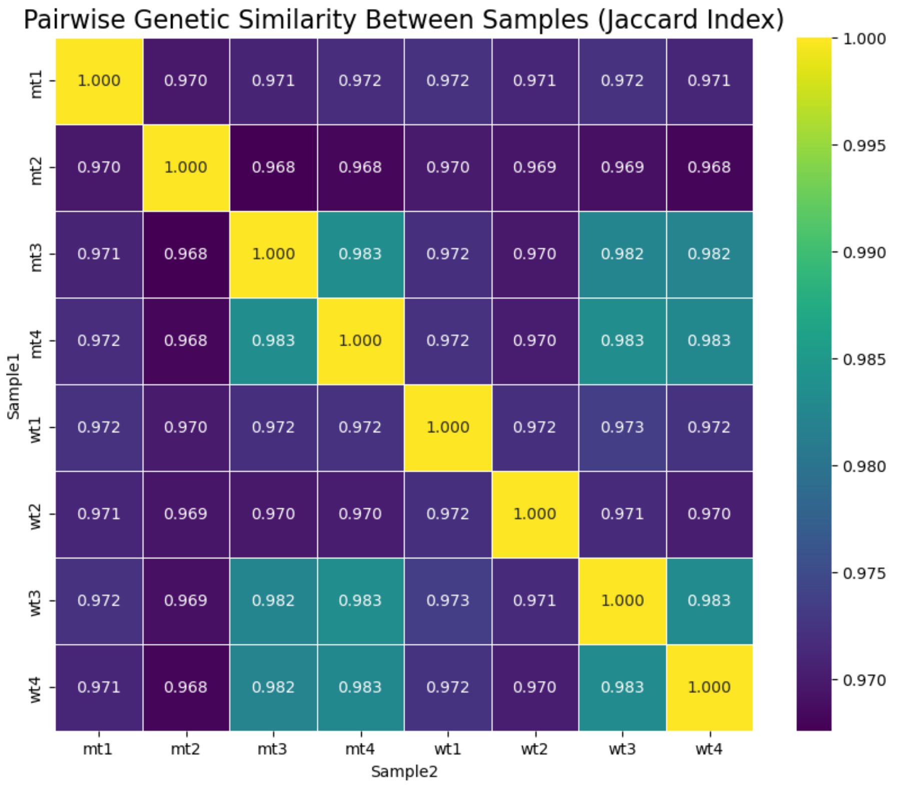


**Figure S1. Heatmap of Pairwise Genetic Similarity Between Samples.** The heatmap visualizes the Jaccard similarity index calculated for all possible pairs of the eight samples. The color scale indicates the degree of similarity, with warmer colors (yellow) representing a higher proportion of shared genetic variants. The clear clustering of mutant (mt1-mt4) and wild-type (wt1-wt4) samples confirms their distinct genetic identity and the integrity of the biological replicates.

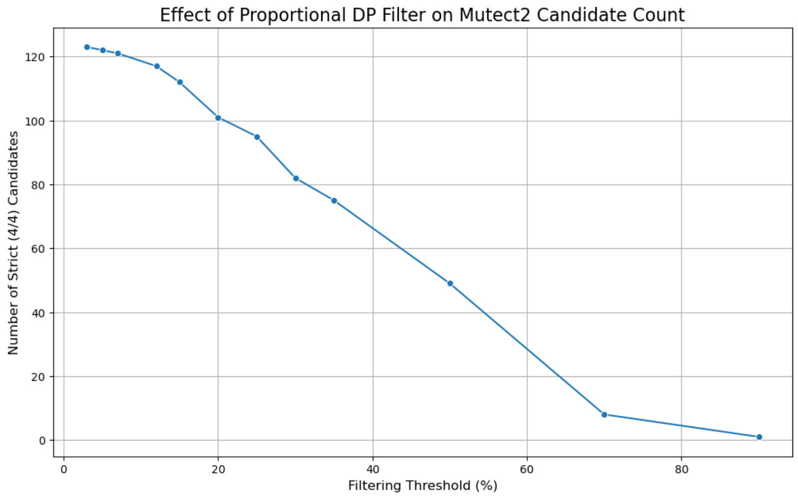


**Figure S2. Effect of the Proportional Depth Filter on Final Candidate Yield from the Mutect2 Pipeline.** The plot shows the total number of high-confidence somatic candidates identified after applying different technical filtering stringencies. The x-axis represents the percentage of mean coverage used as a minimum read depth (DP) threshold, while the y-axis shows the resulting number of candidate variants that pass the strict 4/4 vs. 0/4 biological filter.

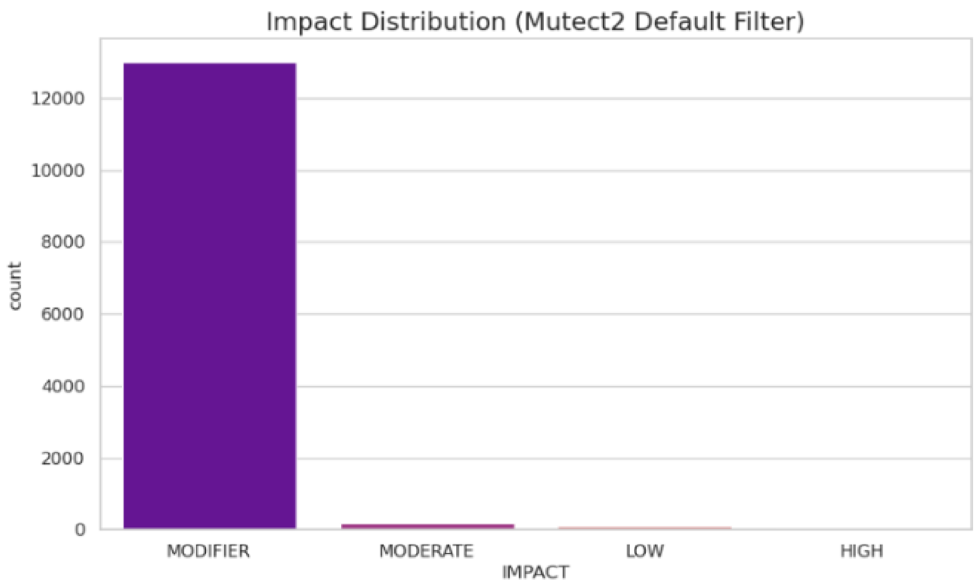


**Figure S3. Impact Distribution of Candidates from the Mutect2 Default Filter.** The plot shows the frequency of each impact category for all candidates identified using only the standard Mutect2 filtering pipeline.

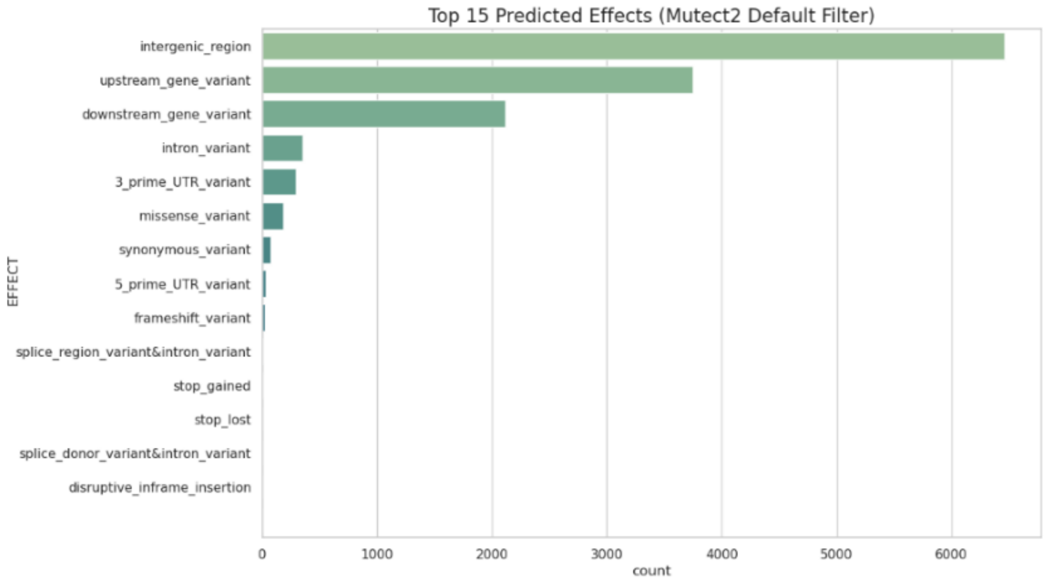


**Figure S4. Top 15 Predicted Effects for the Mutect2 Default Filter Set.** The plot shows the most common variant effects, dominated by those in non-coding regions.

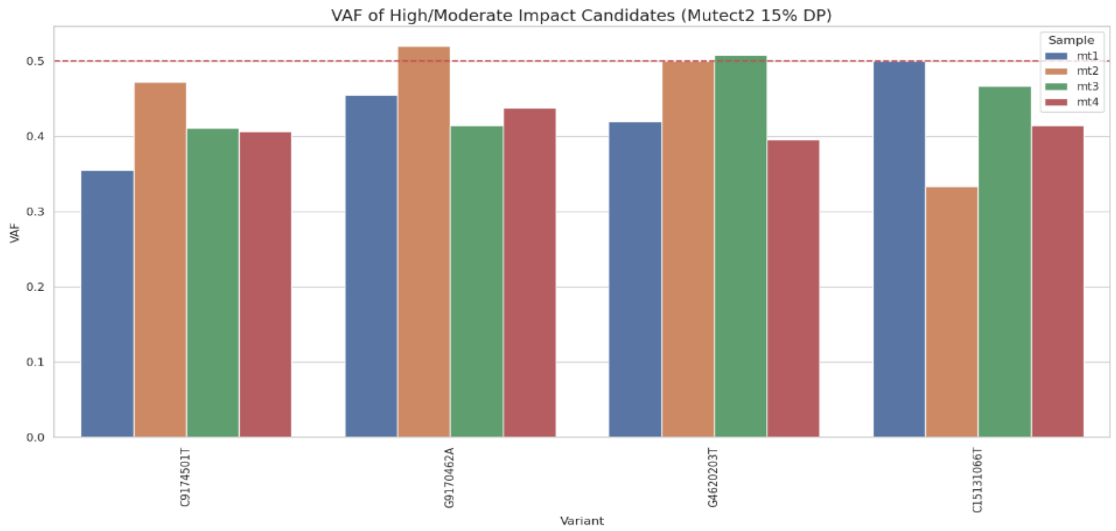


**Figure S5. Variant Allele Frequency (VAF) of Top Mutect2 Candidates.** The plot shows the VAF for high/moderate impact candidates identified after proportional depth filtering. The consistency across all four mutant samples validates these as high-confidence variants.

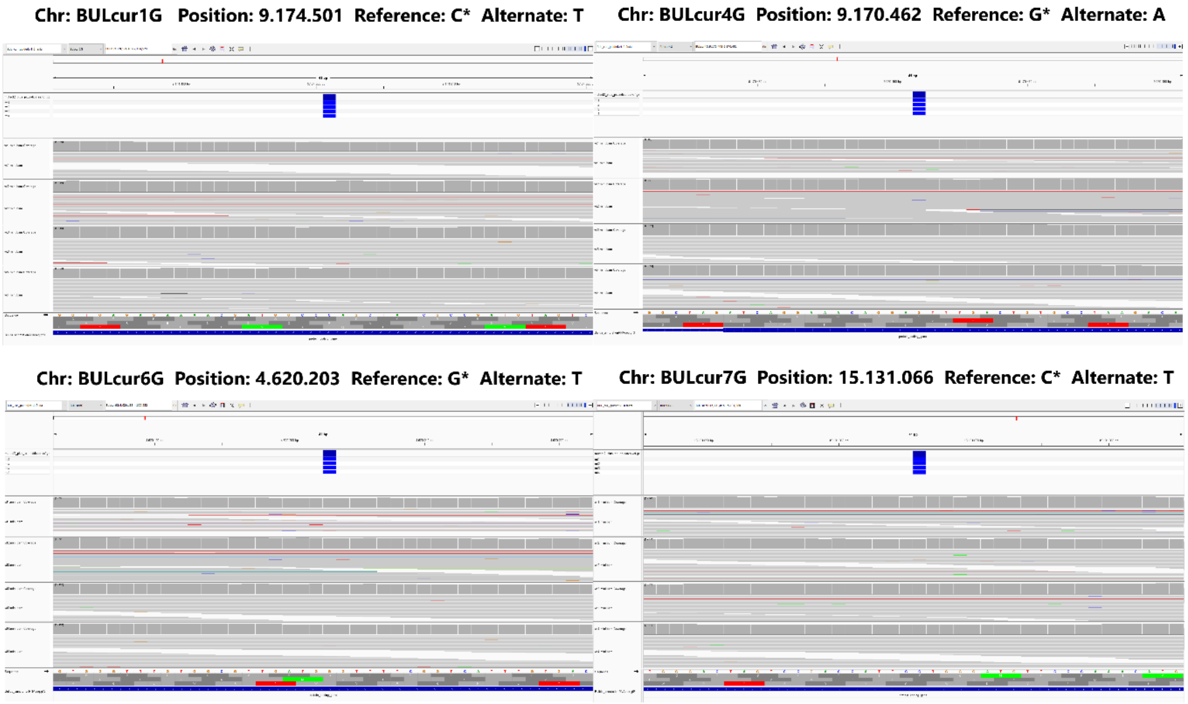


**Figure S6. IGV snapshots of the four final high-confidence candidate somatic mutations.** Each panel shows the read alignments for the four mutant replicates (top tracks) and the four wild-type replicates (bottom tracks) at the indicated genomic position. Non-reference bases are colored, highlighting the alternate allele. (A) A C>T stop_gained mutation in Gene_9712. (B) A G>A missense variant in Gene_26305. (C) A G>T missense variant in Gene_20739. (D) A C>T missense variant in Gene_42548. The consistent presence of the alternate allele exclusively in the mutant samples provides strong visual validation for these candidates.

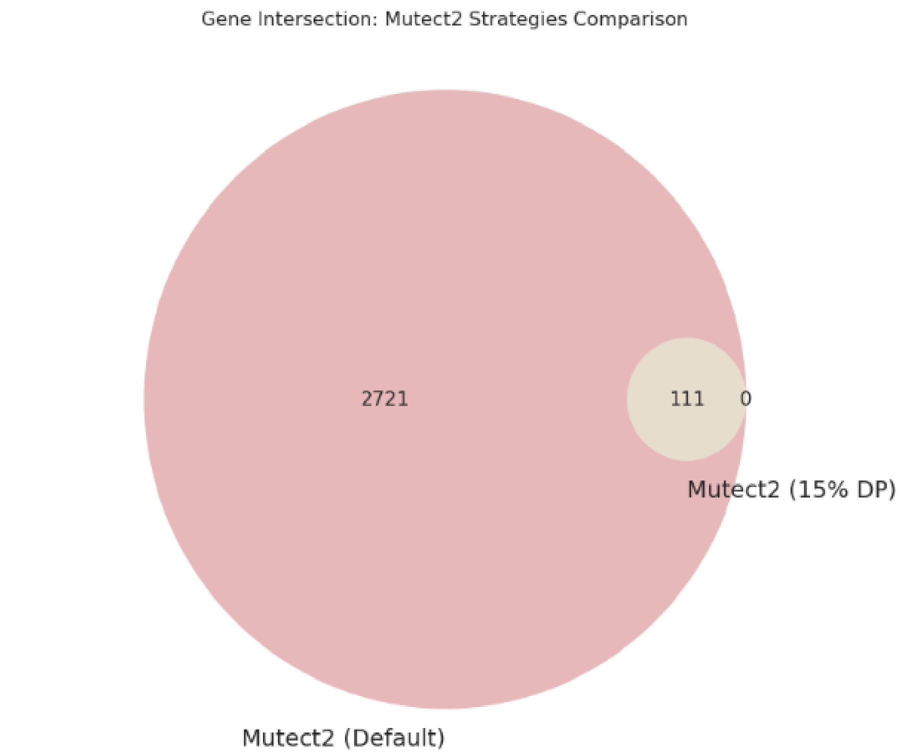


**Figure S7. Comparison of Genes Identified by the Two Mutect2 Filtering Strategies.** The diagram shows the overlap in affected genes between the candidate set from the default filter and the set from the more stringent proportional depth (15% DP) filter.

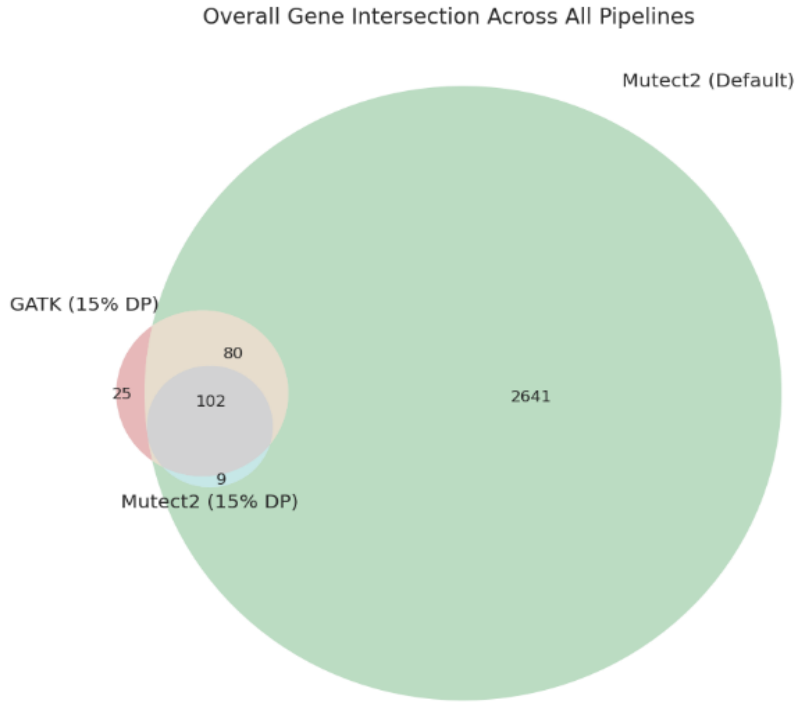


**Figure S8. Three-Way Comparison of Affected Genes Across All Pipelines.** This Venn diagram shows the intersection of the three final candidate gene sets. The central overlap represents the genes identified with the highest possible level of confidence.

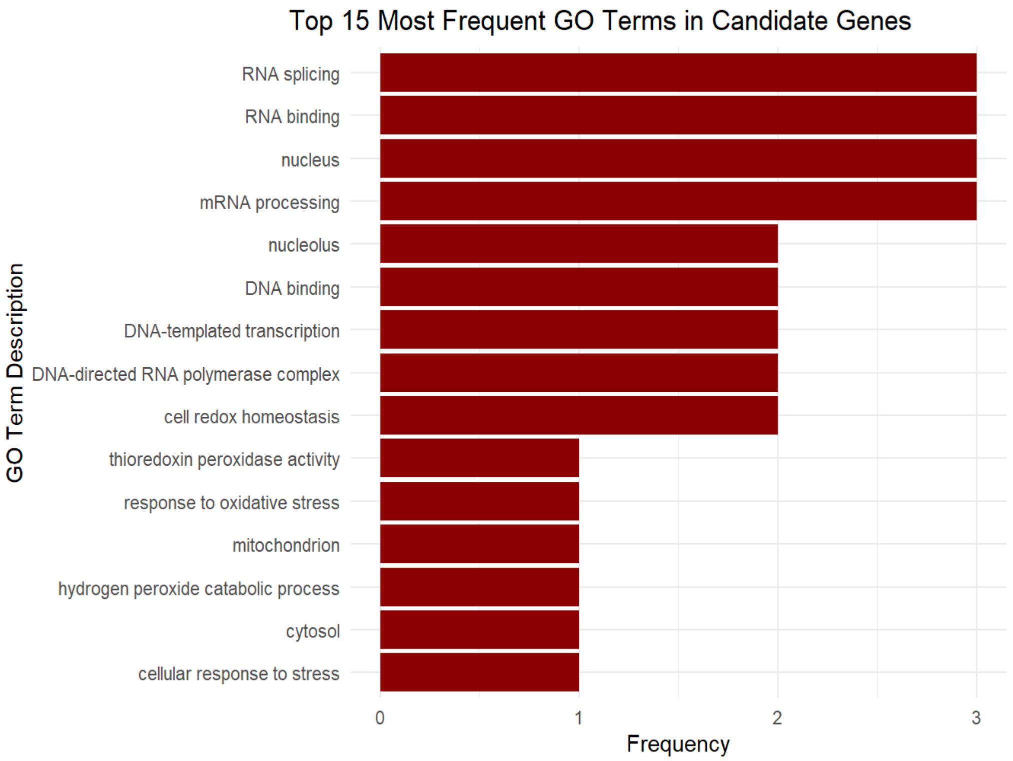


**Figure S9. Frequency of Top 15 GO Terms Among Candidate Genes.** The bar chart displays the count of the most common GO terms associated with the final set of candidate genes.

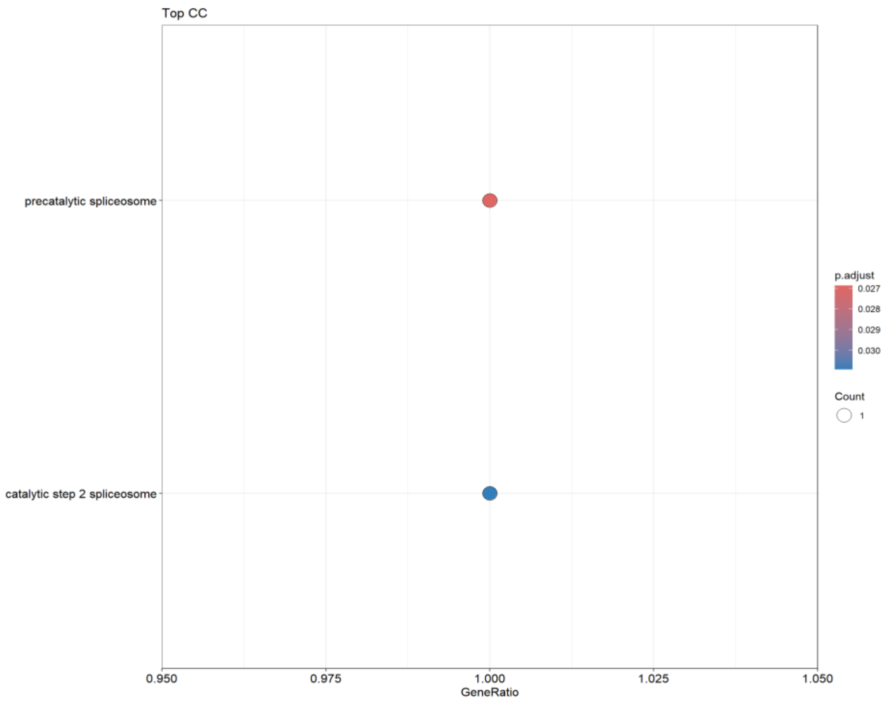


**Figure S10. Enriched Gene Ontology (GO) Terms for the Cellular Component Category.** The dot plot displays the most significantly enriched GO terms. The size of the dot represents the number of genes from the candidate list associated with the term, and the color represents the statistical significance (adjusted p-value).

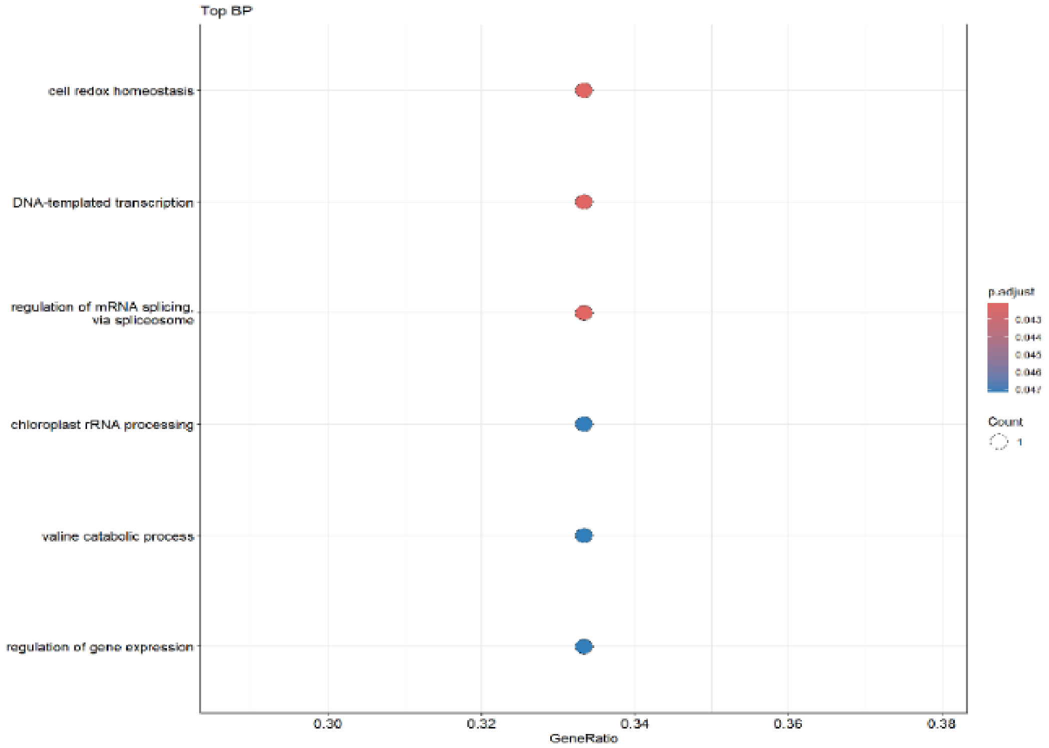


**Figure S11. Enriched Gene Ontology (GO) Terms for the Biological Process Category.** The dot plot displays the most significantly enriched GO terms. The size of the dot represents the number of genes from the candidate list associated with the term, and the color represents the statistical significance (adjusted p-value). The results point towards a significant enrichment in functions related to RNA/DNA processing and gene expression.
